# The role of temporal sensitivity of synaptic plasticity in representation learning

**DOI:** 10.1101/2024.09.27.615415

**Authors:** Patricia Rubisch, Melanie I Stefan, Matthias H Hennig

## Abstract

Synaptic plasticity, the process by which synapses change in an activity-dependent manner, is assumed to be the basis of learning. Experiments demonstrated that synaptic plasticity not only depends on excitatory activity but also on the rate and timing of inhibitory events. Hypothesising that the regulatory effect of inhibition is mediated by membrane potential hyperpolarisation, we identify fast fluctuation sensitivity as a contributing factor to the inhibitory regulation of plasticity. Fast fluctuation sensitivity characterises the influence of fast changes in the membrane potential on the plasticity model predictions, including (short) hyperpolarising events. Furthermore, by introducing a novel plasticity model, the Voltage-Dependent Pathway model, we show that fast fluctuation sensitivity enables a precise temporal regulation of plasticity via inhibition tightly locked to stimulus presentation. In recurrent networks, receptive fields develop independent of the degree of fluctuation sensitivity, yet the temporal precision of inhibitory regulation is critical for the heterogeneity and quality of the resulting representation.

**Author Summary:** Synaptic plasticity plays a central role in learning and enables us to interact within an ever-changing environment flexibly. The term governs all processes which leads to a change in connection strength between two neurons. Previously, studies of synaptic plasticity strongly focused on the role of excitatory activity. However, experiments excluding GABA blockers implicate inhibitory neurons to regulate synaptic plasticity. We further investigate the role of inhibitory neurons as regulators of synaptic plasticity in a model study. Specifically, we focus on the decrease in membrane potential due to inhibitory activity as a key actor in plasticity regulation. This comparative modelling study considers two well-established models and a novel model which differ in the order in which filtering and thresholding is applied to the membrane potential. We find that while all models develop connectivity structures resembling receptive fields characteristics of the visual cortex, the quality and heterogeneity of the developed feature detectors is model-specific. Heterogeneity and representation specificity is promoted by a high degree of fast fluctuation sensitivity of the model enabling temporally precise inhibitory regulation of plasticity determined by the model’s filter and threshold.

## Introduction

Long Term Potentiation (LTP) and Long Term Depression (LTD), forms of activity-dependent plasticity, are the foundational processes by which an organism learns. Standard plasticity protocols describe conditions under which LTP and LTD can be experimentally observed. These include the precise timing between pre- and postsynaptic spikes called Spike-Timing Dependent Plasticity (STDP)^1–3^ and the description of pre-post spike pairing frequency dependency^2,4^ as well as the dependency of LTP/LTD on the presynaptic stimulation frequency^5–7^. Often these experiments are performed in the absence of inhibition^1–3,5,8^ and therefore exclude any effects mediated by inhibitory activity. However, multiple studies show that inhibition influences and changes the established experimental results of the standard protocols^9–13^. Specifically, Hayama et al. ^14^ and Paille et al. ^15^ found that STDP outcomes differ when inhibition in the form of GABAergic activity is present at the synapse. In detail, Hayama et al. ^14^ observed a promotion of LTD when a pre-post spike pair is preceded by inhibition. Moreover, Paille et al. ^15^ demonstrated the reversal of the causal relationship of standard STDP in the presence of inhibition. A direct link between hyperpolarisation due to inhibition and the regulation of plasticity at excitatory synapses is described by Person and Raman ^13^.

Models of synaptic plasticity allow us to explore the functional role of inhibition in plasticity regulation within networks of neurons. In rate- and STDP models inhibition and its influence on plasticity promotes functional development and stability^16–18^. However, they require a-priori assumptions about the relationship between Excitatory-Inhibitory (EI) balance and plasticity updates. We hypothesise that inhibitory regulation effects are already accounted for in voltage-dependent plasticity models avoiding any assumptions on the mechanisms themselves. In voltage-dependent plasticity models, plasticity induction is dependent on the relationship between the membrane potential and (arbitrary) plasticity thresholds. Thus, any hyperpolarisation of the postsynaptic membrane potential caused by inhibitory activity directly impacts the plasticity updates. Therefore, they provide the opportunity to investigate inhibitory regulation of plasticity without specifying the interaction between inhibition and the plasticity update explicitly. Moreover, a detailed study of inhibitory regulation in voltage-dependent plasticity models is currently missing.

We address this gap by introducing a novel plasticity model, the Voltage-Dependent Pathway model (VDP), and by systematically comparing it against a well-established model introduced by Clopath et al. ^19^ (CM) and its variation as presented by Meissner-Bernard et al. ^20^ (MB) with regard to their sensitivity to fast membrane potential fluctuation. We find that the fluctuation sensitivity determines the effect of inhibition through the extent to which hyperpolarising events contribute to plasticity updates. The higher fluctuation sensitivity of the VDP enables a tighter temporal regulation of plasticity by inhibition. We show that this drives synaptic development linked to input and inhibition onset. Overall, high sensitivity to fast fluctuation of the membrane potential promotes receptive field heterogeneity improving the overall quality of the developed input representation.

## Results

In voltage-dependent plasticity models, we assume that the direction of plasticity updates depends on the postsynaptic membrane potential. Thus, the models are not only sensitive to postsynaptic spike timing, but also predict plasticity updates in the absence of postsynaptic spikes^19–21^. In the following we assume that inhibition affects plasticity updates through its hyperpolarising effect on the membrane potential, and we investigate this effect in voltage-dependent plasticity models.

### Hyperpolarisation is sufficient to model inhibitory regulation of STDP

Voltage-dependent plasticity models are based on electrophysiological experiments demonstrating that the sign of the plasticity update is related to the depolarisation level of the synaptic membrane^8,22–30^. Although well-established models have demonstrated voltage-dependent plasticity models widen the range of reproducible experimental results^20,21^, they only consider standard plasticity protocols, during which inhibitory signalling is often blocked. However, Paille et al. ^15^ and Hayama et al. ^14^ report that GABAergic signalling within a typical STDP pairing changes the plasticity outcome. We used an STDP protocol with precisely timed inhibitory inputs to assess the relationship between membrane potential, inhibitory timing and predicted plasticity updates. Here, we compare three models of this type: The model presented in^19^(CM), which is the foundational voltage-dependent model, its variation introduced in^20^(MB) and a novel model called the Voltage-Dependent Pathway model (VDP). All of them implement Hebbian plasticity where the sign and amplitude of the update are dependent on the difference between the postsynaptic trace(s) and the plasticity thresholds *θ*^+,−^. Based on their formulation (summarised in Table 1, mathematical details provided in Methods), they can be divided into two subclasses: filter-first-threshold-second (MB, CM) and threshold-first-filter-second (VDP).

**Table 1:** Summary of the treatment of the pre- and postsynaptic activity by the different voltage-dependent models in the LTP and LTD induction conditions. All models assume individual thresholds for LTP and LTD induction for which *θ*^+^ *> θ*^−^.

| Model | update | presynaptic | postsynaptic |
| --- | --- | --- | --- |
| MB<sup>20</sup> | LTP & LTD | low-pass filtered | threshold on low-pass filtered |
| CM<sup>19</sup> | LTP | low-pass filtered | threshold on low-pass filtered & instantaneous |
|  | LTD | spike time (instantaneous) | low-pass filtered |
| VDP | LTP & LTD | low-pass filtered | threshold on instantaneous followed by down-stream dynamics |

In STDP simulations, including a precisely timed inhibitory event relative to the postsynaptic event, the effect of filtering the membrane potential before thresholding in the MB and CM becomes apparent. Filtering the membrane potential results in a prolonged effect of the inhibitory event. Even inhibitory activity delivered after the postsynaptic spike influences the STDP kernel and shifts its LTP-LTD ratio, see Figure 1B and C. In comparison, the effective time range in the VDP is restricted to negative inhibition-post intervals.

**Figure 1.**
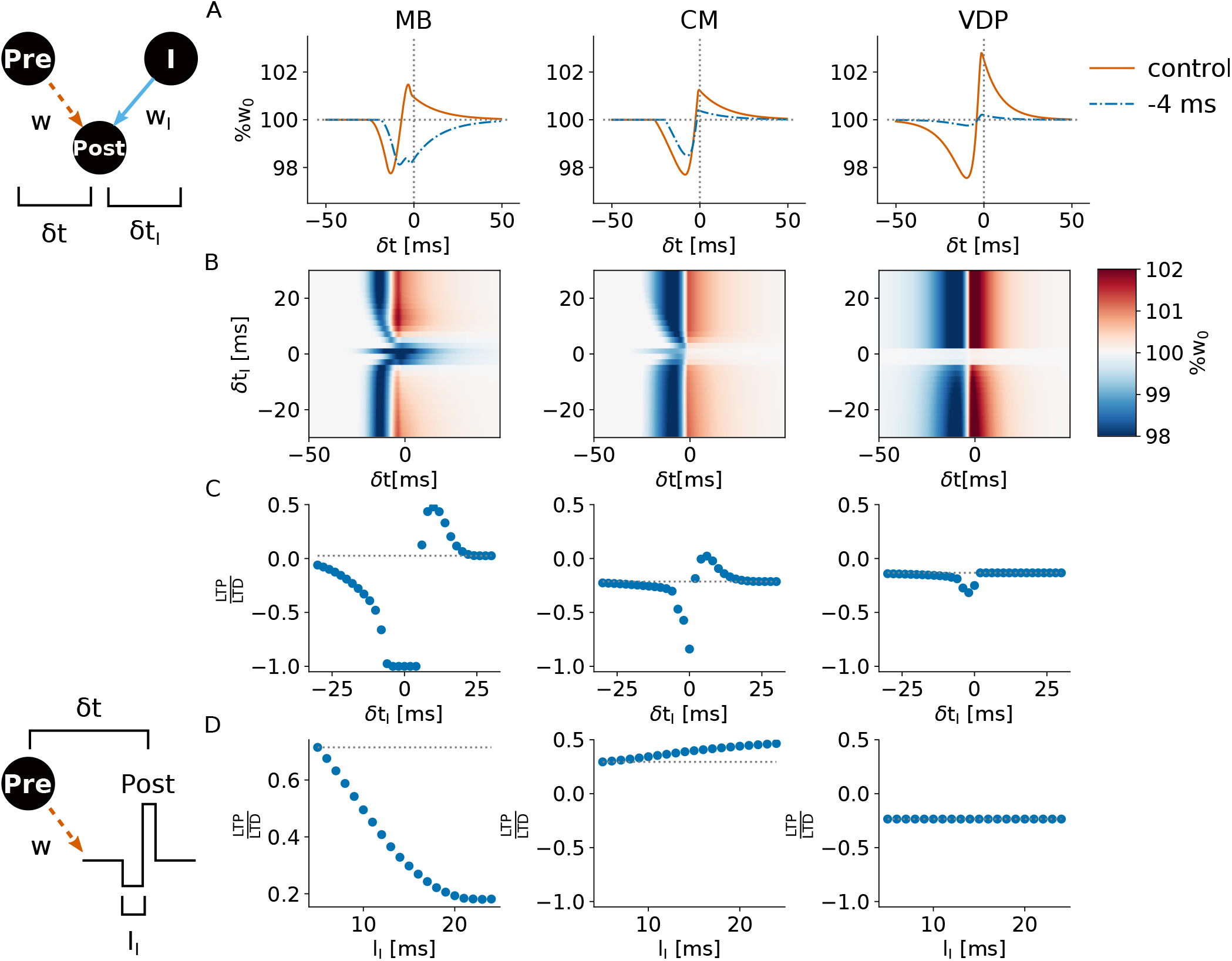
Inhibition mediated via hyperpolarisation in voltage-dependent plasticity models regulates synaptic change in standard STDP experiments. A standard STDP protocol is extended by an inhibitory presynaptic neuron as indicated by the circuit diagram. The inhibitory spike time *δt*_*I*_ is measured relative to the postsynaptic spike. Models are indicated by the abbreviations on top. **A:** Example STDP kernels of the standard protocol without inhibition (orange) and with inhibitory spiking at *δt*_*I*_ = −4ms. **B**: STDP kernels over several different inhibitory spike timings shown as a heatmap. Blue areas indicate LTD, while red denotes LTP. The MB and CM allow inhibition to affect the plasticity update over a broader range of inhibitory spike timings. **C**: LTP-LTD ratio of the kernels shown in **C**. Pre-inhib-post pairings lead to promotion of LTD. Pre-post-inhib pairings lead to promotion of LTP. **D**: Inhibitory STDP protocol but the inhibitory spike is replaced by a hyperpolarisation of length *I*_*I*_ as shown in the visualisation to the left. *δ*_*t*_ denotes the interval between the presynaptic spike and postsynaptic depolarisation. Filter-first models show a length-dependency, while the VDP’s plasticity results are independent of the length of inhibitory event.

Paille et al. ^15^ reported that the LTP reverts to LTD for pre-post pairings. The MB model is the only model out of the three which shows such a polarity change of the plasticity update. Pre-inhib-post pairings induce LTD while the pre-post pairings without inhibition result in LTP. In the CM and VDP, the inhibitory activity decreases the amplitude of both LTP and LTD. However, computed the LTP-LTD ratio reveals that LTP is more strongly attenuated. Overall, the LTP-LTD ratio shifts towards LTD-dominance (Figure 1C).

Another difference between the filter-first-threshold-second models and the VDP is the effect of the hyperpolarisation duration(Figure 1D). Both the MB and CM are sensitive to the length of the inhibitory event. The longer the duration, the stronger the shift of the LTP-LTD ratio. The ceiling of the effect is caused by the leaky integration of the neuron model and the filtering of the membrane potential by the plasticity models. In contrast, the VDP is insensitive to the duration of the inhibitory stimulus. Only the strength of the hyperpolarisation influences the plasticity update. In summary, the inhibitory STDP experiment offers first insights into the differences between the models with regard to inhibitory regulation of plasticity. In filter-first threshold-second models inhibition has a broader effective window and larger effect amplitude. A model which applies the thresholds first, followed by downstream dynamics, like the VDP, allows only for inhibitory activity preceding the postsynaptic spike to effect synaptic plasticity disregarding of the duration of the hyperpolarisation.

### The order of thresholding and filtering determines fast fluctuation sensitivity of the model

The results above illustrate how plasticity predictions for isolated spike pairs are affected by inhibition and differ between the models. These differences are caused by the order of filtering and thresholding in the model definitions. Low-pass filtering dampens fast components of the signal. Thus, filtering first reduces the sensitivity to fast fluctuations by removing them. To investigate fast fluctuation sensitivity in the models, we replace the postsynaptic spikes of the previous protocol with a fluctuating membrane potential, where we manipulate the fluctuation speed and strength.

To simulate the postsynaptic membrane potential of a neuron receiving input from many neurons, we simulate it as an Ornstein-Uhlenbeck (OU) noise. It is defined by its fluctuation speed *τ* and standard deviation *σ* which determines the amplitude of the fluctuations (see example traces in legend of Figure 2A).

**Figure 2.**
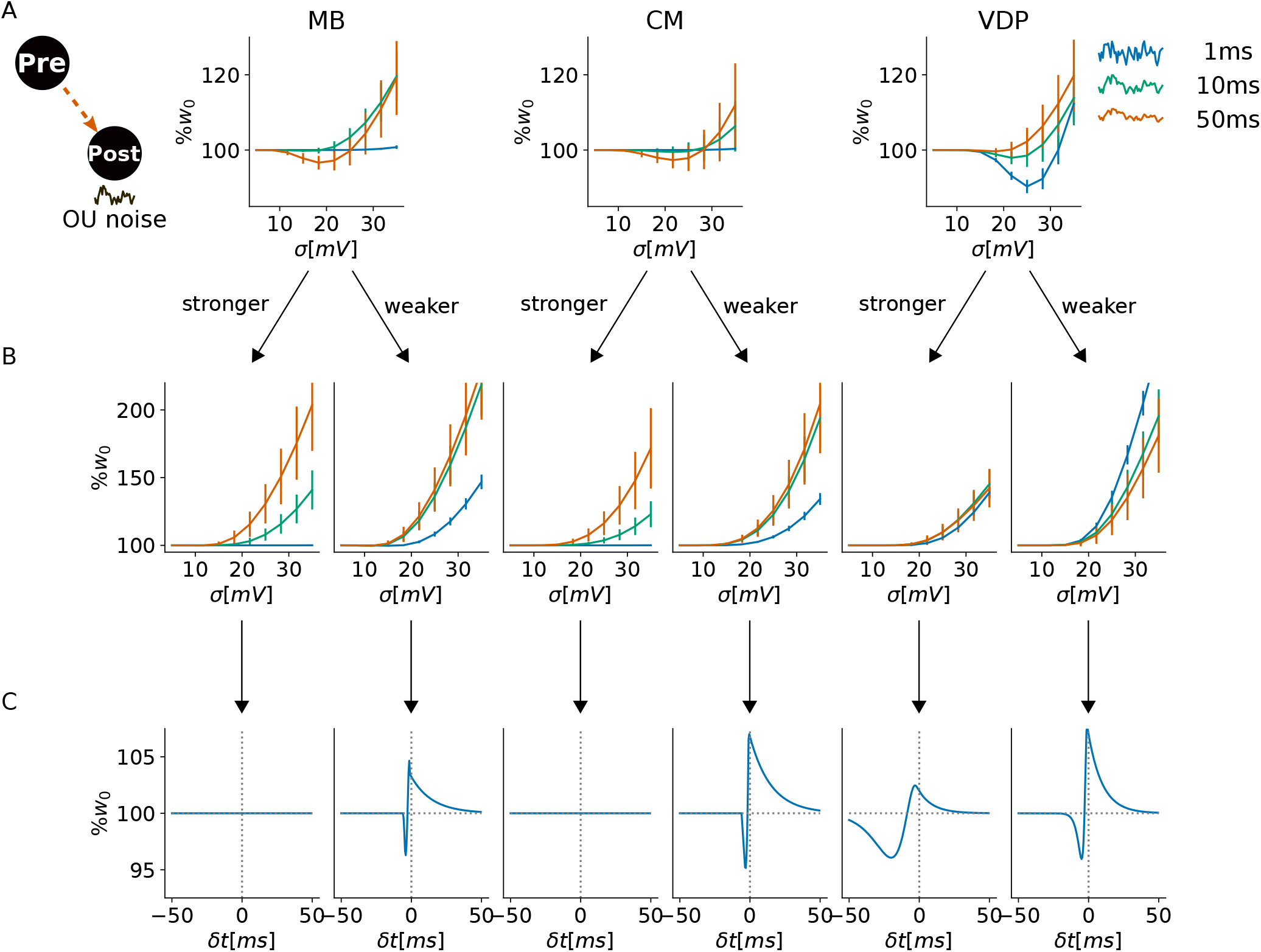
Fast fluctuation sensitivity is a model feature of the VDP while the established models show a parameter-dependent sensitivity. **A:** The postsynaptic membrane potential *v* is replaced by an Ornstein-Uhlenbeck (OU) noise. Results are obtained over a 10s simulation and averaged across 100 instantiations. To control the fluctuation speed, the time constant *τ* of the OU noise is varied (colours) while increasing the standard deviation *σ* results in growing fluctuation amplitude. Plasticity models are indicated by their abbreviations as in previous figures. **B:** Parameter changes which increase filter strength (“stronger”, synaptic time constants increased) or decrease filter strength (“weaker”, synaptic time constants decreased) demonstrate that fast fluctuation sensitivity can be achieved in the MB and CM with a weak filter. However, the VDP’s fast fluctuation sensitivity is independent of parametrisation. **C:** Corresponding STDP kernels to the parameter setting above. Strong filters abolish STDP kernels by reducing the magnitude of the filtered signal to below threshold values. Weaker kernels reduce the LTD window in all models. In the MB and CM, LTD induction is reduced to near instantaneous pairings, while the VDP model retains a wider window in comparison.

The results confirm that weight changes in the VDP model have a higher sensitivity to fast fluctuations than the MB and CM (Fig. 2)A. Filtering reduces high-frequency components, which are further attenuated by thresholding the filtered signal. In contrast, thresholding first preserves the overall frequency structure of the original signal (see Supplementary for illustrations). As the MB and CM are filter-first-threshold-second models, they show a reduced sensitivity to fast fluctuations. Therefore, their plasticity updates are driven by slow and high-amplitude events.

These observations are not unique to the parameters used in these simulations. The high sensitivity to fast fluctuations of the VDP plasticity updates is preserved for a range of parameter variation (Fig. 2B). Both the MB and CM also show an increase in fast fluctuation sensitivity when the postsynaptic filter time constant is reduced. Reducing the filter time constant, decreases the time window over which plasticity occurs shrinking STDP considerably (Fig. 2C). In the VDP, the LTD window also shrinks, but to a much lesser extent than in the CM and MB models.

Increasing the postsynaptic filter constants leads to further reduction in sensitivity to fast fluctuations for the MB and CM, and the models no longer show STDP kernels. These can be recovered by reducing the threshold of both LTP and LTD induction as spikes are strongly attenuated due to the strong filtering. In contrast, the VDP model predicts STDP kernels despite increasing the postsynaptic filter constants. The filter constants do not affect the membrane potential or plasticity threshold. Therefore, the the temporal relationship between membrane potential threshold crossings and presynaptic spikes is preserved accross parameter variation in the VDP. However, the downstream dynamics are either prolonged (“stronger”) or shortened (“weakened”) which leads to an wider/shorted LTD induction window.

To recap, filter-first-threshold-second models more strongly attenuate the postsynaptic signals, which reduces the sensitivity to fast fluctuations and results in a parameter-dependent sensitivity. In contrast, the fast fluctuation sensitivity of the VDP is a model feature, preserving realistic STDP kernels across postsynaptic filter strengths across parameters.

### Slow averaged signals drive rate-dependency

The simulations above demonstrate how single spikes and fast signals are picked up by the different plasticity models. Next we analyse the relationship between fluctuation sensitivity and plasticity at different average firing rates. Experiments have shown that LTD is induced at low frequencies while the synapse strengthens at high presynaptic frequencies^5–7^. To simulate biologically realistic postsynaptic activity, a neuron receives activity from an excitatory and inhibitory Poisson population (Fig. 3A). This allows us to not only examine the frequency dependency of plasticity, but also the influence of the EI balance of the synaptic input.

**Figure 3.**
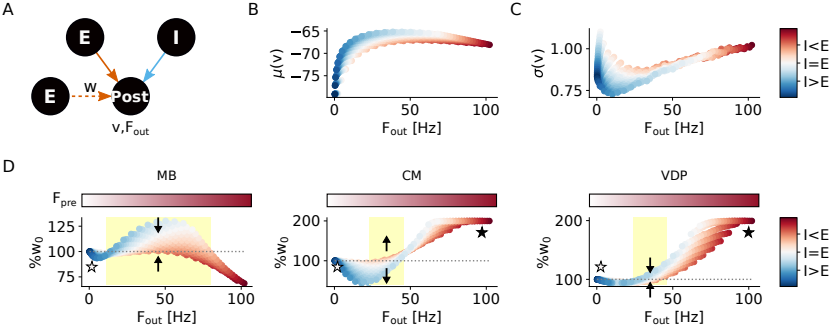
The nonlinearity of frequency-dependency is driven by slow and averaged signals, resulting in small differences between the plasticity models. EI balance variation at constant postsynaptic frequency can shift LTP to LTD and vice versa. **A** A postsynaptic neuron is driven by 100 excitatory and 100 inhibitory Poisson neurons with firing rates *F*_*E*_, *F*_*I*_ ∈ [0, 100)Hz (5Hz resolution). The colour-coding of the data points indicates the EI balance of the input computed as 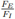 with 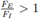 coloured red and 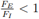 coloured blue. We measure the postsynaptic membrane potential, output frequency, as well as the weight change of an excitatory (orange,dashed) synapse. The presynaptic neuron of that synapse fires regularly with the same frequency as the excitatory Poisson population. All results are calculated over 10s simulation time. **B**: The mean membrane potential increases when the synaptic input is inhibition-dominated. **C**: The standard deviation of the membrane potential measuring the fluctuation amplitude is increasing when the synaptic input is dominated by excitation. **D**: Plasticity predictions of the plasticity models of the excitatory weights. Model identity is indicated by the abbreviations on top. All results are averaged over 100 runs. The color bar on top indicates the frequency of the regular spiking presynaptic neuron. The open-faced star indicates the regime of pathological behaviour at low postsynaptic frequencies (negative plasticity-activity feedback). The black star indicates the regime of pathological behaviour at high postsynaptic frequencies (positive plasticity-activity feedback). Yellow-backed regions indicate intermediate frequency prediction of special interest. Arrows indicate the direction of EI balance shift due to the predicted synaptic plasticity update.

In voltage-dependent models, postsynaptic frequency is an indirect factor in determining the plasticity update. Prolonged depolarisations which do not generate a spike can induce plasticity. Therefore, the mean membrane potential and its variance is more informative than postsynaptic frequency alone. While counter-intuitive, the mean membrane potential (Fig. 3B) is higher when the synaptic input is dominated by inhibition. Simultaneously, the standard deviation decreases with stronger inhibition at the same postsynaptic frequency (3C). These statistics are similar across neuron models and not limited to the Leaky-Integrate-and-Fire model used here (see Appendix). It is important to note that the standard deviation does not measure fluctuation speed, but the fluctuation amplitude around the mean membrane potential.

The interactions between frequency-dependency and EI balance (Fig. 3D) are driven by the differences in membrane potential statistics. At similar frequencies, inhibition-dominated synaptic input results in a higher mean membrane potential compared to the postsynaptic activity elicited by excitation-dominated synaptic input. For models which are sensitive to the mean of the membrane potential, this causes LTP to be more strongly promoted by inhibition-dominated synaptic input. However, this is not true for all three models. The CM model predicts predominantly LTD for inhibitory-dominated synaptic input (*I > E*) and LTP for excitatory-driven activity (*I < E*). In contrast, the MB and the VDP predict LTP for synaptic input characterised as *I > E* and LTD for excitatory-dominated input. For an excitatory synapse, LTP updates increase the EI balance while LTD updates decrease it, pushing it further towards *I > E*. Thus, the CM progressively pushes excitatory synapses to the extreme points: to the minimal weight for inhibitory-dominated input and to the maximal weight for excitatory-dominated input. In contrast, the MB and VDP push excitatory synapses towards their starting value.

However, the CM and VDP models show similarities at the extreme condition where inhibitory activity is low. Both models demonstrate a positive feedback between excitatory weight update and postsynaptic frequency. This positive feedback results in exploding weights and has been studied as a feature of Hebbian plasticity^31^.

Overall, all models predict a non-linear relationship between the postsynaptic frequency and the resulting plasticity updates. However, investigating the effect of EI balance variation of the synaptic input demonstrates qualitative different behaviour between the models.

### Fast fluctuation sensitivity enables tight inhibitory control during learning of structured input

The previous simulations focused on scenarios where postsynaptic neurons either emits single spikes or are driven by constant activity. However, real-world statistics of neural activity are time-dependent and change systematically with different stimuli. Synaptic plasticity allows neurons to learn these patterns, with both single spikes and rates influencing synaptic development.

In the following, we aim to gain an understanding of the interaction between the rate-dependenc STDP effects and difference in fluctuation sensitivity during learning of structured input. We specifically focus on timed inhibitory signalling to investigate inhibitory regulation during stimulus changes.

Three postsynaptic neurons are driven by an input population which is separated into three distinct groups. At each point in time two out of the three input groups are active. Additional input to each postsynaptic neuron is provided by a dedicated inhibitory population. The activity pattern of the input groups and the inhibitory population are inverse of each other: When input population 1 and 2 are active and population 3 is silent, inhibitory population 3 is active while 1 and 2 are in-active. We are interested in the development in the input synapses in order to investigate how the synaptic development is driven by the timed inhibition. In order to differentiate we divide the synapses into to groups: The encoding weights and the non-encoding weights. The encoding weights connect the input population with its corresponding postsynaptic neuron (E1 to 1, E2 to 2, E3 to 3), while the non-encoding weights include the remaining synapses (E2/E3 to 1, E1/E3 to 2 and E1/E2 to 3). Figure 4A and B show an illustration of the set-up and activity.

**Figure 4.**
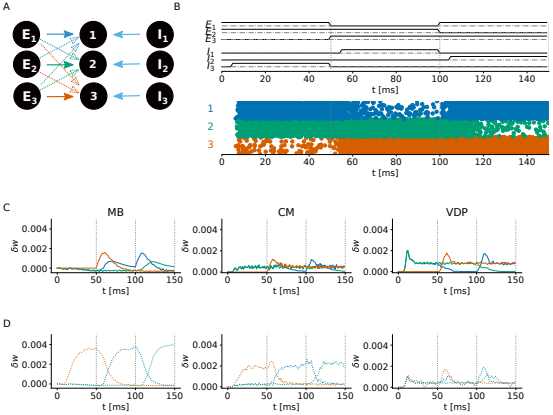
Sensitivity to fast fluctuations promotes stimulus-locked plasticity updates. The higher temporal precision of the inhibitory regulation increases the VDP’s sensitivity and reaction speed to changes in the received synaptic input. **A**: Three distinct excitatory Poisson neuron populations (N=10) drive neural activity of 3 LIF neurons. Each LIF neuron receives inhibition at a predefined time from its dedicated inhibitory Poisson neuron population (N=10). Solid arrows indicate encoding weights which connect a specific Poisson population with the neuron which is inhibited when the input population is in-active. Dotted lines indicate non-encoding weights. All results are means of 100 independent runs. **B**: Postsynaptic firing rate of the excitatory neurons and the activation pattern of the input and inhibitory populations. **C** Synaptic update over time of the encoding weights. **D**: Synaptic updates of the non-encoding weights.

We consider two key factors when comparing the weight development between plasticity models: the amplitude difference between encoding and non-encoding weight change and the temporal pattern of the update predictions. The update size of non-encoding and encoding weights differ strongly between the models. The MB’s and CM’s maximal non-encoding synaptic change is significantly larger than the synaptic change of the encoding weights. In contrast, the results of the VDP model show similar amplitudes for both the non-encoding and encoding synapses.

The temporal pattern of the encoding weights are fairly similar between the models. The weight updates reaches their maximum shortly after the stimulus changes and the first presentation does not show any updates for the encoding weights of neuron 3 due to no presynaptic activity. However, the non-encoding weight updates show strong differences. The MB and CM weight changes ramp up towards a plateau over a presentation period. In contrast, the VDP’s weight change can be described as a short-timed bump activity. It occurs after the stimulus changes, but the second half of the presentation period is characterised by a constant size of the weight changes. While the time point at which the maximal amplitude is reached is fairly similar for the development of the encoding weights between models, the development of the non-encoding weights differ visibly. The MB model results reach their peak synaptic change at the end of the stimulus presentation. It slowly increases over the presentation time window. The CM model results show a prolonged plateau of at the maximal weight change. This plateau is quickly reached. Therefore, the synaptic update size is constant for half the duration of a single stimulus presentation. In contrast, to the ramping behaviour of the CM and MB model, the VDP updates reach their maximum shortly after the stimulus changes which is followed by a decrease in update size. Similarly to the CM results, the VDP results also show a constant weight update size in the second half of the presentation window.

The higher amplitude of weight updates of non-encoding synapses as well as the ramping behaviour demonstrate that synaptic development in the MB and CM model is strongly driven by slow signals. The weight update increases when the neuron receives inhibition, which is driven by the increase in the mean membrane potential. Visualising the instantaneous activity of the postsynaptic neurons, results in a similar, superposable pattern. In contrast, the VDP weight updates resembles an indicator of change in the postsynaptic activity rather than the development of the activity itself. Weight updates are maximal when the postsynaptic activity changes, but decrease to a constant level quickly after.

The results presented here demonstrate the importance of the fast fluctuation sensitivity during learning of structured input. The MB weights are driven by slow signals due to both LTP and LTD induction relying on filtered signals. In the CM, LTP is sensitive to the instantaneous membrane potential which drives synaptic development after presentation changes. However, LTD is still driven by the filtered membrane potential, a slow signals. Only the VDP, a model in which LTP and LTD take into account the instantaneous membrane potential shows synaptic development tied to sudden changes in postsynaptic activity driven by inhibition onset. Overall, this highlights the importance of fast fluctuation sensitivity of the plasticity to enable a tighter temporal link between activity changes due to inhibition and weight development.

### High fluctuation sensitivity promotes heterogeneity of learned representations

The results above demonstrate the fluctuation sensitivity and prevalence of fast and/or slow signals in determining the plasticity outcomes predicted by the different models. Based on these observations, we presume that the sensitivity to fast signals also plays a pivotal role in the development of representations in recurrent neural networks. In the following, we use two established experiments, the Bars-and-Stripes problem^32^ and receptive field development from natural input statistics^33,34^, to investigate differences in representation learning between the models and connect the developed representations to to well-known properties of the visual cortex.

Brito and Gerstner ^35^ demonstrated that non-linear Hebbian plasticity implements a general principle for receptive field development and unified the biologically motivated plasticity models with principled approaches such as Independent Component Analysis (ICA). The non-linearity specifically refers to the frequency dependency aka the dependence on slow signals, a characteristic all models demonstrate. Thus, they should be able to perform ICA as well.

To test this, networks are presented with the bars-and-stripes problem or Földiak’s task^32,36^. This problem consists of randomly sampled bars and stripes which are presented at the same time, thus forming crosses (Fig. 5A). A network which successfully performs ICA on this input develops receptive fields resembling the independent components, a bar or a stripe, and not the presented crosses. As the non-linear frequency-dependency of all models suggest, every network develops single component receptive fields. However, the results vary in their overall statistics and characteristics of non-single component receptive fields (Fig. 5B-C). The MB network develops predominantly multi-factor cells, showing not only one bar or stripe but up to four independent factors encoded by a single excitatory neuron. Interestingly, these cells do not necessarily resemble the presented input but might consist of two stripes instead of a cross. The CM network is the only network which develops “copying” cells. These cells do not show any discernible pattern in their receptive fields. All their synapses reach the maximal weight causing the cell to always be active. It also develops neurons with opposite connectivity patter: uniform synapses with 0 weights. Neurons with this connectivity are non-functional, as their activity is not driven by feed-forward input but only recurrent activity. The VDP network also develops such 0 receptive fields.

**Figure 5.**
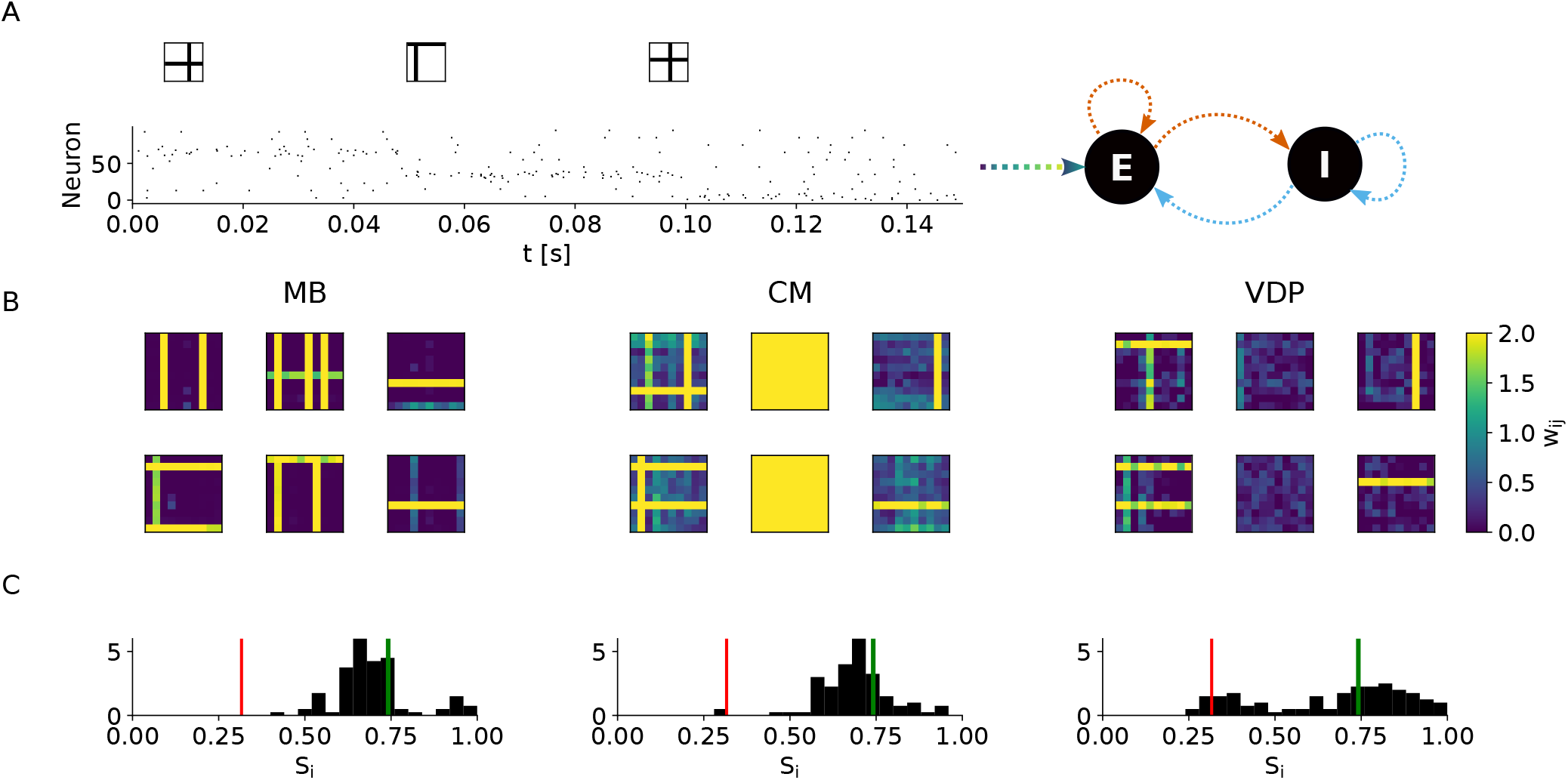
The overlap of learned representations of independent components is reduced in plasticity models with high fast fluctuation sensitivity. **A**: An EI network is presented with the bars-and-stripes task^32^. **B**: Learned receptive fields. **C**: Separation scores of the excitatory neurons. A neuron which encodes a single bar perfectly has *S*_*i*_ = 1.0. The green line indicates the upper bound for a multi-factor cell. The red line indicates the corresponding lower bound. Any score below this characterises random receptive fields with no discernible pattern. These scores are computed with cleaned-up receptive fields to reduce the contribution of spurious patterns present in near-0 weights. All weights *w*_*ij*_ *< µ*(*w*) are set to 0. The mean is computed over all receptive fields of the network.

The separation score *S*_*i*_ measures the quality and quantity of the receptive fields (Fig. 5C). Based on its definition, we can define lower bounds for single-factor receptive fields (green line) and multi-factor cells (red). Single-factor cells can either encode a bar or a stripe. Their score is the same. The histograms plotted in Figure 5C show that the VDP develops the highest number of single-factor cells and overall heterogeneous receptive fields. The separation score distribution is bimodal with one peak at single-factor scores and a second one at the transition from random receptive field scores to multi-factor cells. The tight temporal window of the inhibitory regulation in the VDP results in a higher degree of representational separability. This confirms our presumption that tight temporal inhibitory regulation promotes functional development in recurrent networks. Furthermore, the development of 0-weight synapses is a form of pruning and shows hallmarks of sparse coding. The network consists of more neurons than the input has factors. Thus, not all neurons are needed to represent the input.

The bars-and-stripes problem illustrates that the variation in fluctuation sensitivity drives differences between learned representations. However, as an experiment it is far removed from biological relevant input. When the concept of ICA is applied to naturalistic input, it results in oriented edge-detectors similar to receptive fields in V1^33–35,37,38^. Therefore, we present the MB, CM and, VDP network with CIFAR10 input patches^39^ in order to learn receptive fields resembling those in the visual cortex. In the following we investigate whether the variation in fluctuation sensitivity also promotes differences in the quality and heterogeneity of receptive fields learned from naturalistic input.

The CIFAR-10 input patches are presented by two separate input populations, an On- and Off-population, similar to On- and Off-cells in the retina. Neurons in the On-population fire proportionally to the positive part of the presented image while the value of the negative pixels scales the Off-neuron’s firing rate (Fig. 6A). The receptive field of an excitatory neuron is equal to the difference between its Off-weights and its On-weights. It is important to note that homogeneous receptive fields with *w*_*i*_ = 0∀*i* such as those present in the MB network (see Fig. 6B) do not necessarily indicate all synapses to be silent as in the previous experiments. While this can be the cause, perfectly aligned, equally strong On-and Off-synapses cancel out also resulting in a receptive field with 0 weights.

**Figure 6.**
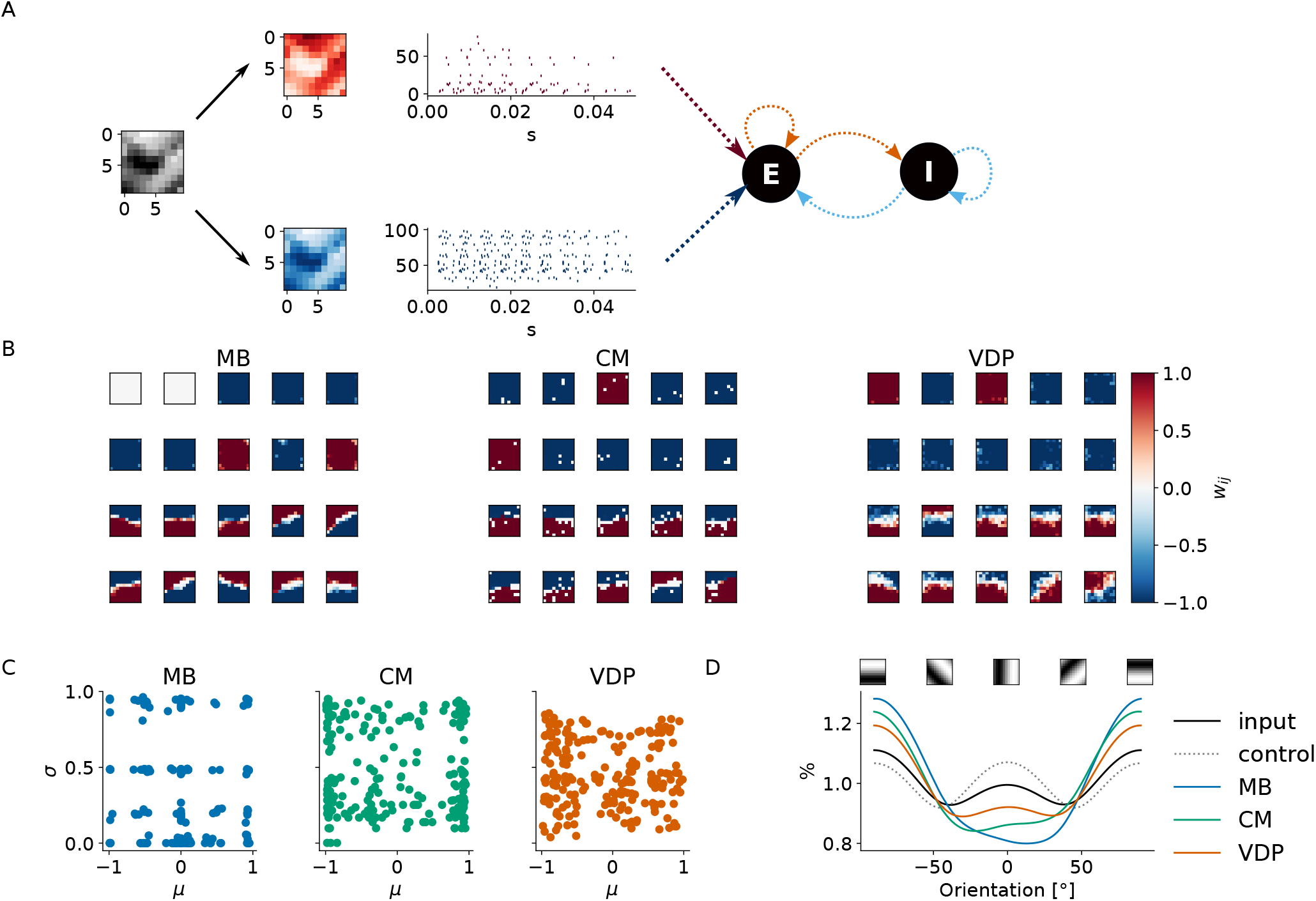
Receptive field quality and heterogeneity are promoted by plasticity models with high sensitivity to fast signals. **A**: A fully-plastic recurrent EI network is presented with CIFAR10 input patches using an On- and Off-population for input encoding. **B**: The developed receptive fields show specific characteristics depending on the plasticity model. Shown are 10 receptive fields with the smallest standard deviation and 10 receptive fields with the highest standard deviation. Blue indicates a strong connection to the Off-population, while red indicates a strong synapses to the On-population. **C**: Mean and standard deviation of the receptive fields. **D**: Responsiveness of oriented Gabor-patches to the input data (black), random patches (grey-dotted), receptive fields of the MB (blue), CM (green) and VDP-network (orange). The colour scheme for the receptive fields is consistent between **C** and **D**.

All networks develop a mixture of (uniform) mono-polar and oriented receptive fields (Fig. 6B). Mono-polar receptive fields are characterised by a small variance and the mean *µ* is either −1 or 1 (Fig. 6C). These receptive fields copy the input from either the On- or Off-population. The MB network receptive fields show the strongest clustering of their statistics (*µ* and *σ*) in comparison to the others. They are dominated by mono-polar and 0-mean-receptive fields. The CM receptive fields also shows some clustering at *µ* = [1, −1] in their statistics. However the standard deviation scores are covering the value range more uniformly, leading to less clear clusters compared to the MB statistics. The statistics of the VDP receptive fields show even less clustering. This suggest that the VDP network developed the most diverse representations while the MB receptive fields include a lot overlapping patterns.

The other characteristic which is of interest when talking about receptive fields is their orientation. The visual cortex has several orientation columns in which neurons are sensitive to the same orientation and can detect edges in images^40^. To compute the orientation, the receptive fields are convolved with predefined Gabor patches or varying orientation between −180° and 180°. The relative response patterns as shown in Figure 6D indicates how closely the different network fit the distribution of orientations of the input data.

Horizontal edges are dominant in the input data. In comparison to diagonal orientations, vertical features show a stronger response, but not to the same extent as horizontal orientations. Overall, this leads to a “w”-shape of the orientation profile of the input data where the bump of the w is lower than its tips. This distinct “w”-shape is only present in the response profile of the VDP network receptive fields. However, the difference between the response contribution of horizontal and vertical edges is bigger than compared to the input data response profile. Neither the MB nor the CM profile can be desribed as a “w”-shape. The, the middle bump is absent in both. This indicates that neither network learned vertical orientations well. Interestingly, the MB receptive fields underrepresent orientations with a small positive angle and show the highest relative response to horizontal edges. In contrast, edges with small negative angles are less present in the receptive fields developed by the CM. Furthermore, the difference between the response to horizontal stimuli and vertical/diagonal oriented gabor patches is bigger compared to the VDP response and the input data. Therefore, the receptive fields developed by the VDP represent the input data the closest. Even though, the absolute contributions do not exactly match, the overall shape of the response profile is well approximated.

In summary, the characteristics of the developed connectivity structures in these two functional experiments suggest that the higher sensitivity to fast signals of the VDP enabling a tight temporal inhibitory plasticity regulation, promotes the separability of input representations. The receptive fields show a higher degree of heterogeneity and resemble the orientation biases present in the input data more closely. Therefore, the degree of fast fluctuation sensitivity directly affects the quality of the developed representations in a recurrent network.

## Discussion

These results suggest that the inhibitory regulation through hyperpolarisation of the membrane potential is a plausible model for developing functional connectivity patterns. A detailed analysis of three voltage-dependent plasticity models highlights the importance of the sensitivity to fast signals to enable precise inhibitory regulation. The stronger the filtering of the postsynaptic membrane potential, the weaker the sensitivity to fast fluctuations. In recurrent networks, the sensitivity to fast fluctuation is important to enable accurate input representation due to inhibitory signalling.

Biologically, these results suggest that synapses located on distal dendrites can rely more on the local, fast-changing information^41–43^, if plasticity has higher temporal sensitivity, such as proposed in the VDP. This implies that distal synapses might develop more precise and distinct representations, as they are less influenced by longer temporal integration. In contrast, at proximal synapses, the voltage signal corresponds to information integrated over longer periods of time and over many synapses. For this case our simulations suggest plasticity will yield representations with more overlaps. As the models by Clopath et al. ^19^ and Meissner-Bernard et al. ^20^ do not have a high temporal sensitivity, they are predicted to behave similarly at distal and proximal synapses. In contrast, the VDP predicts a difference in representation learning at distal and proximal locations. This prediction is testable in biophysical neuron models, and differences in distal and proximal tuning properties should be detectable in imaging experiments.

Neurons and synapses are highly heterogeneous: from genetic markers to protein and channel sub-type composition and local distribution^43–45^. Thus it is unlikely that a single rule can account for all behaviours. Here, we only considered synapses reproducing the standard STDP kernel, while several different STDP kernel forms have been identified experimentally^1^. These models cannot reproduce effects such as the the reversal from LTD to LTP described by Paille et al. ^15^, as it is incompatible with voltage-dependent plasticity models where the LTD threshold is always below the LTP threshold^8^.

Interestingly, all models show a non-linear frequency-dependency. While we can argue that the frequency-dependency is driven by slow signals such as the mean membrane potential, the models also differ in their level of sensitivity to this slow signal. This is detectable in the reported effect of the synaptic EI balance driving the postsynaptic frequency. We want to stress, that while the experiment approximates a mean-field analysis of the model’s behaviour, it is not sufficient to make claims about the stability of the models. A detailed exploration of the activity development over long simulations of recurrent networks, including parameter sweeps, would be necessary for an accurate investigation of model stability. Unfortunately, this is outside the scope of this work which focuses on the role of model characteristics on representation learning.

Similarly, the models implement Hebbian plasticity without considering homeostatic components. Homeostatic plasticity increases network stability and can be implemented by a plethora of mechanisms^31^. Thus, future research investigating network stability in voltage-dependent plasticity rules should also include a detailed exploration of the interactions with homeostatic principles acting on biologically plausible time scales.

This work exclusively focuses on the inhibitory regulation in voltage-dependent plasticity models. While this family of plasticity models is well positioned to investigate hyperpolarisation as a mean of inhibitory regulation, it is not the only one possible to address this. Calcium-dependent plasticity models can also incorporate a direct link between the membrane potential and calcium influx and are able to reproduce experimental data incorporating neural architecture^46,47^. Specifically, Hiratani and Fukai ^47^ demonstrates the development of dendritic EI balance in their work. However, the work is not investigating the effects of representation learning. Furthermore, the insights derived here and the categorisation of models can be extended to them. Both models would be classified as filter-first-threshold-second and therefore would show a decreased sensitivity to fast signal components. What these components might signal and their contribution to the activity of downstream processes in biological processes remains to be further investigated. In summary, this work investigates the role of inhibition in voltage-dependent plasticity models. Inhibitory regulation of plasticity in these models is determined by their sensitivity to fast fluctuations. The higher the fast fluctuation sensitivity, the tighter the temporal inhibitory regulation of plasticity. By studying representation learning and stimulus/inhibition-onset sensitivity, we identified that a high sensitivity to fast signals is important for recurrent inhibition to effectively drive plasticity and functional development. The novel plasticity model, the VDP, demonstrates that a high degree of fast fluctuation sensitivity results in a more heterogeneous receptive fields which accurately represent the input data. This work predicts that synapses whose plasticity is driven by local, fast-fluctuating information like those on distal dendrites develop more distinct and precise input representations due to the precisely timed inhibitory plasticity regulation. Investigating the relationship and interplay between local EI balance and LTP/LTD induction at different dendritic sites warrants further investigations to accurately describe the functional role of inhibitory regulation of plasticity in biological neurons.

## Methods

### Neuron model

All experiments, except the STDP experiments, are performed using a Leaky Integrate-and-Fire (LIF) neuron with conductance-based synapses. They follow the equation:

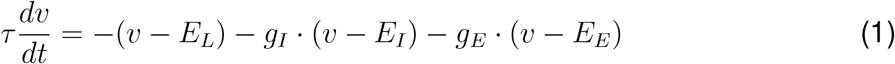

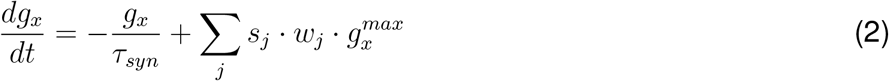

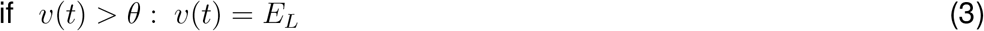

with *x* = [*E, I*] and *E*_*L*_ denoting the resting potential of the neuron. *g*_*I*_ and *g*_*E*_ are the synaptic conductances of the inhibitory and excitatory synapses. Whenever a spike arrives at the synapse *g*_*x*_ grows proportionally to the current synaptic weight *w*_*x*_. We use 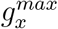 to denote the maximal conductance. We generally assume the notation of *w*_*ij*_ denoting the synapse connection between presynaptic neuron *j* and postsynaptic neuron *i*. Therefore, *j* in Equation 2 indicates that the synaptic current is summed over all presynaptic neurons. The standard parametrisation of the LIF is: *E*_*L*_ = 79*mV, E*_*E*_ = 0*mV, E*_*I*_ = −80*mV* and *τ* = *τ*_*E*_ = *τ*_*I*_ = 10*ms*.

In order to enable a fair comparison of isolated events as occurring in STDP experiments at low repetition frequencies, the results presented in Figure 1 and Figure 2C use a Generalised LIF (GLIF) model of class 5 as introduced in Teeter et al. ^48^. The following equations define this model:

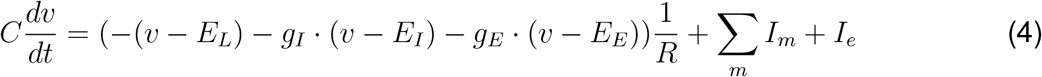

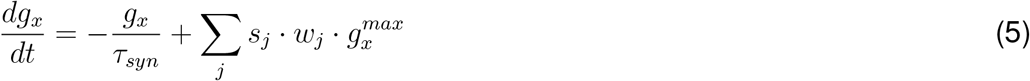

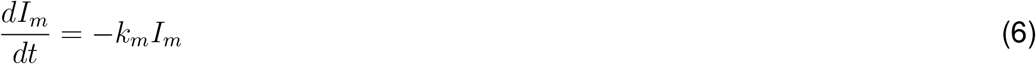

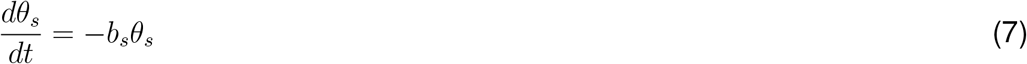

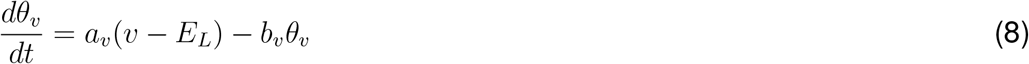

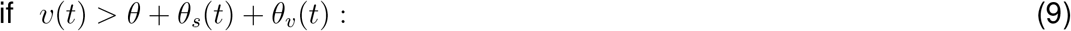

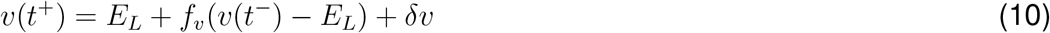

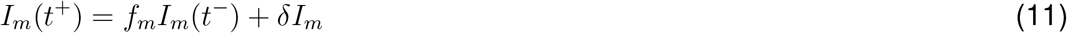

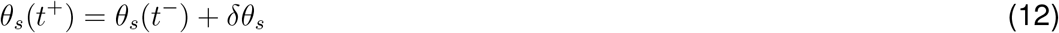

*t*^+^ denotes the time step after spike generation, while *t*^−^ denotes the timestep before spike generation. The synaptic conductance *g*_*x*_ follows the same dynamics as in the LIF model. Teeter et al. ^48^ include a full description of the parameters and variables, including explanations of biological equivalence. The parameter setting used here is simulating a neuron of cluster 1^48^, producing excitatory neuronal behaviour. *I*_*e*_ denotes the external current applied to force a spike at precise times.

The GLIF5 neuron model includes After-Spike-Currents (Eq.6) which approximate the spike shape. Thus, the depolarisation is prolonged in the GLIF5 model after spike generation in comparison to a LIF neuron model. This enables models which use postsynaptic membrane filters before thresholding (MB and CM), to predict updates in STDP experiments despite isolated postsynaptic events. The GLIF5 model is therefore similar to the AdaptiveExponential (AdEx) neuron model which was used by Clopath et al. ^19^ to introduce the CM.

### Voltage-dependent plasticity models

All three voltage-dependent plasticity models implement Hebbian plasticity. They can be divided into their presynaptic component and their postsynaptic component. The presynaptic component follows the same dynamics in all models:

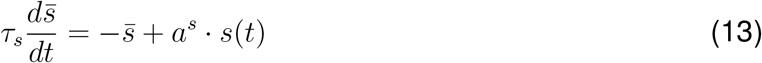

The models either use the presynaptic spike train *s*(*t*) or the presynaptic trace 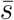. The spike train *s*(*t*) is 1 if a spike was generated at time *t* and is 0 otherwise. 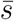 is computed by applying a low-pass filter to *s*(*t*). The scaling parameter *a*^*s*^ is set to 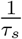 if not indicated otherwise.

A further commonality between the models is the use of an explicit potentiation and depression threshold, *θ*^+^ and *θ*^−^ respectively. *θ*^+^ *> θ*^−^ is true for all parameter settings used in this work. This assumption follows the results presented in Ngezahayo et al. ^8^.

#### Voltage-Dependent Pathway model

The Voltage-Dependent Pathway (VDP) model is a novel plasticity model and can be summarised as a threshold-first-filter-second model. The membrane potential is first passed through a sigmoidal function *σ*(*v*(*t*) − *θ, k*) with the plasticity threshold *θ* and k = 0.4. The outcome sets the update amplitude of the pathway activity *p*(*t*). The pathway activity implements the postsynaptic filter whose properties are demonstrated in the Appendix. It is defined as bi-exponential differential equations:

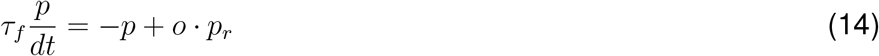

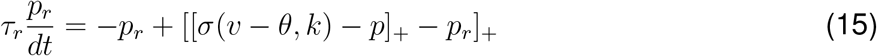

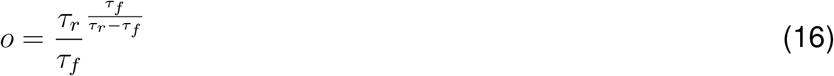

where [*·*]_+_ denotes a linear rectifier: 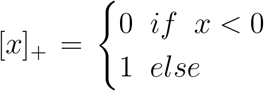 The plasticity update is computed by the difference between the potentiation and depression pathway, *p*^+^ and *p*^−^ respectively. We omitted the superscripts ^+,−^ in the definition for the pathway activity *p* above to improve legibility. This means that all parameters in the bi-exponential definition are pathway specific. Therefore, the dynamics of the potentiation pathway *p*^+^ is determined by the values of 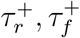 and *θ*^+^. The difference between the potentiation and depression pathway dynamics is multiplied by the presynaptic trace to compute the overall plasticity update. This ensures that updates only occur if presynaptic and postsynaptic activity coincides within a specified time window determined by the pathway specific parameters.

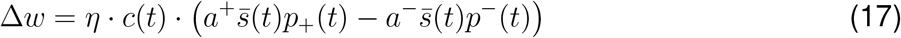

*η* denotes the learning rate, while *a*^+^ and *a*^−^ are pathway specific amplitudes. *c*(*t*) is a scaling factor which introduces a dependence on the external calcium concentration *Ca*^2+^ and is based on the results reported in Inglebert et al. ^4^. This factor is dependent on both the activity of 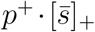 and *p*^−^:

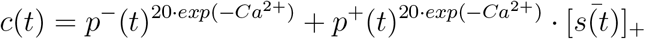

This uncommon formulation causes a stronger expression of depression at low calcium levels. However, the results presented here are independent of this scaling factor because it does not effect the fluctuation sensitivity of the plasticity model. A step-by-step example of the induction mechanism of the VDP is illustrated in the Appendix. As standard parametrisation the VDP was fit to a standard STDP kernel^3^ (see Appendix). We refer to the values of time constants and threshold as the standard parametrisation as these determine the dynamics of the model. These are 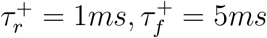, *θ*^+^ = −40.75*mV*, 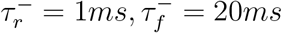, *θ*^−^ = −43.75*mV* and *τ*_*s*_ = 10*ms*. The values of scaling factors are provided in the Appendix which lists all used parametrisations and their corresponding results.

#### The Clopath et al. ^19^ and Meissner-Bernard et al. ^20^ models

The CM and MB model are in contrast filter-first-threshold-second models. The membrane potential *v* is low-pass filtered to compute the synaptic trace 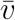. 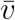 is than directly compared against the plasticity thresholds *θ* in the weight update equation. Potentiation and depression are dependent on their respective postsynaptic traces 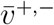 and plasticity thresholds *θ*^+,−^.

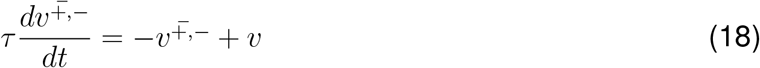

The difference between the CM and MB model lies in the exact combination of the postsynaptic traces and the presynaptic trace. The CM defines the weight-update as follows:

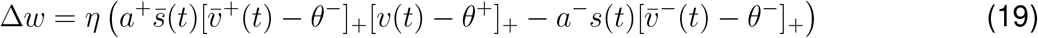

It uses the spike train *s*(*t*) for the induction of LTD. Thus, depression can only occur at the time of a presynaptic spike. Potentiation is conditional on the instantaneous membrane potential. Therefore, both the membrane potential as well as the postsynaptic potentiation trace need to be above a certain threshold.

In comparison the MB model updates synaptic weights following this definition:

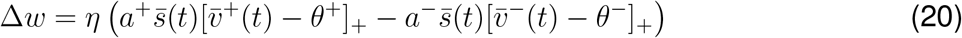

Updates only occur depending on the pre- and postsynaptic traces. Thus, any synaptic weight change is determined by filtered signals, decreasing the contribution of fast signals.

Due to the similar definition of the models, their standard parametrisations are equivalent. They follow the parametrisation described in Clopath et al. ^19^ (thresholds have been adapted due to difference in neuron model): *τ* ^+^ = 7*ms, τ* ^−^ = 10*ms* and *τ*_*s*_ = 15*ms*. A full list of all parameters per experiment is given in the Appendix.

### Simulation Set-up

All simulations are carried out using the Euler integration method with an integration time step of 0.1*ms*

#### Spike-Timing-Dependent-Plasticity with Inhibition

The synapse of interest is the connection between a regular spiking presynaptic neuron (*F* = 0.1*Hz*) and postsynaptic neuron. The postsynaptic neuron is simulated using the GLIF5 neuron model with an input current of *I*_*e*_ = 5*nA* which is applied for 0.8*ms*. The input current is delivered with a frequency of *F* = 0.1*Hz* with a spike time interval *δt* relative to the presynaptic neurons spike. *δt* takes values in the range of −50*ms* to 50*ms* with a spacing of 1*ms*. To include inhibition in the STDP protocol, the postsynaptic neuron also received input from an inhibitory neuron, whose spikes are timed relative to the postsynaptic neuron. This timing difference is denoted as *δt*_*I*_ and ranges from −20*ms* to 5*ms* (resolution of 1*ms*).

To test the effects of the length of an inhibitory event at constant peak membrane potentials, we approximate the postsynaptic hyperpolarisation and spike as a step-wise membrane. Inhibitory events hyperpolarise the membrane potential by 5mV for a duration of *I*_*I*_*ms* and followed by a depolarisation to *E*_*L*_ + 30*mV* for 1*ms*.

The LTP-LTD ratio *r* is computed via:

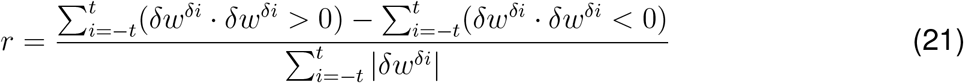

It computes the absolute sum of all positive and negative weight updates over all measured *δt*, subtract them from each other and scales it by the sum of all absolute weight changes. Therefore, *r* is restricted to be between −1 and 1.

#### Fast fluctuation sensitivity using Ornstein-Uhlenbeck membrane potential dynamics

To investigate the varying degree of fluctuation sensitivity of the plasticity models (Figure 2), the postsynaptic membrane potential is simulated as Ornstein-Uhlenbeck noise. It follows the dynamics governed by:

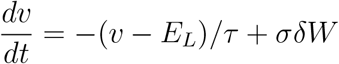

*δW* is a Wiener process. The resting potential *E*_*L*_ is equivalent to the resting potential of the LIF neurons of other experiments with *E*_*L*_ = −79*mV*. Conditions include a variation of the fluctuation speed *τ* = [1, 10, 50]*ms* and amplitude *σ* = (5, 35)*mV*. Presynaptic activity is simulated using a regular spiking neuron with frequency *F* = 10*Hz*. Each condition is simulated for 10*s* with 100 repetitions. The presented results are averages over the repetitions computed for each condition defined by *τ* and *σ*.

#### Frequency-dependency

To approximate the frequency dependency, two separate Poisson populations drive the postsynaptic LIF neuron. The simulation includes an explicit excitatory and inhibitory presynaptic population to account for the variation in sub-threshold membrane potential dependent on the synaptic input composition. Their activity is set by *F*_*E*_ and *F*_*I*_ which take values between 1 and 100*Hz*. The synaptic EI balance which is visualised by the colour of each data point is defined as the ratio 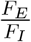. The presynaptic neuron which connects the plastic synapse to the postsynaptic neuron fires regularly with the same frequency as their corresponding Poisson population. Thus, the presynaptic frequency in Figure 3D is *F*_*E*_. Results showing the inhibitory synaptic development with presynaptic frequency *F*_*I*_ are illustrated in the Appendix. The simulation is run for 10*s*. Each condition is repeated 100 times and results are averages computed across repetitions.

#### Inhibition/Input-Onset specificity of synaptic updates

To investigate the effect of rapidly changing input, three postsynaptic LIF neurons receive activity from an input population(*N*_*input*_ = 30) and an inhibitory populations (*N*_*I*_ = 30). To structure the network, the input population is divided into three groups based on their activity and the inhibitory population is subdivided by their activity and synaptic connections. The input groups are all-to-all connected to the postsynaptic neurons. In contrast, each of the three inhibitory sub-population is connected to one dedicated postsynaptic neuron. Both inhibitory and input population consists of Poisson neurons. When active, the input neurons’ firing rate is set to *F*_*input*_ = 100*Hz* and the inhibitory neurons fire with a rate of *F*_*I*_ = 50*Hz*. The activity pattern during one presentation is the following: Two out of the three input-groups are active. The inhibitory sub-population of the inactive input-population inhibits their corresponding postsynaptic neuron. Therefore, when E1 and E2 are active, I3 is active (see Figure 4A). Each presentation lasts for 50*ms* and each possible pattern is presented once. Results are shown as the synaptic updates averaged over 100 instantiations of this simulation.

#### Bars-and-Stripes

The Bars-and-Stripes experiment of Földiak’s bars^32^ consists of bars and stripes which are simultaneously presented. Each pixel of the 10×10 pixel input patches is represented by a Poisson neuron in the input popualtion. When the pixel is part of the presented overlapping strip and bar, the neurons firing rate is set to *F*_*input*_ = 100*Hz*. This input is presented to a fully-connected recurrent network with *N* = 120 (*N*_*I*_ = 0.2 *· N*_*E*_). The initial input weights are all-to-all connected to the excitatory network. All synapses are plastic.

To assess the quality of the developed representations, we compute a separation score *S*_*i*_ based on the receptive field *r*_*i*_ of each excitatory neuron *i*. The receptive field *r*_*i*_ is defined as all weights *w*_*ij*_ which connect the postsynaptic (excitatory neuron *i*) with the input population. Therefore, *j* denotes the presynaptic neuron for which*j* ∈ *N*_*input*_. The separation score *S*_*i*_ measures the maximal cosine-similarity *c*_*S*_ between the receptive field *r*_*i*_ and the ideal receptive fields of all bars 0° and the ideal receptive fields of all stripes 90°:

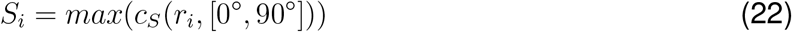

Prior to computing *S*_*i*_, we clean the receptive fields *r*_*i*_ by discarding any weights which are smaller than the overall average postsynaptic connection. This decreases the contribution of residual patterns in non-functional, small synapses. When the receptive field encodes a single factor the cosine similarity *c*_*S*_ to this factor is 1.0, while the similarity to all remaining factors is smaller than 1. The presence of multiple factors in the receptive fields, disregarding whether in the form of a cross or two non-intersecting factors, decreases the cosine similarities between the receptive field and any given single factor. Based on this, we can compute an upper bound for multi-factor receptive fields, which simultaneously provides a lower bound for a single-factor cells (*S*_*i*_ = 0.74). Additionally, we compute an upper bound for non-informative random fields (*S*_*i*_ = 0.32).

#### Receptive field development from naturalistic input

1000 Patches from the CIFAR10 data set are presented to a fully recurrent EI network with *N* = 250, *N*_*I*_ = 0.2*·N*_*E*_. Each of the centered 10×10 pixel input image (*µ* = 0) is encoded by two dedicated neurons. One neuron is part of the On-population and the other belongs to the Off-population. Neurons in the On-population are active relative to the positive of the image patch, while the Off-population neuron’s frequency is set according to the negative of the image. The neurons in either population follow the dynamics of:

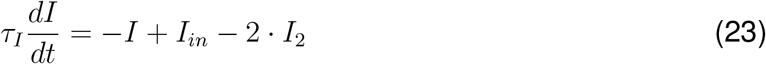

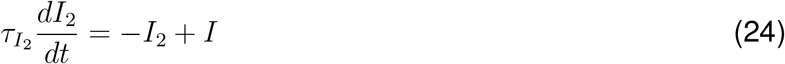

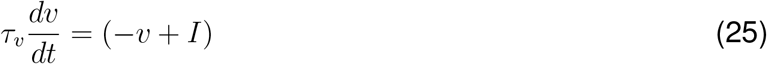

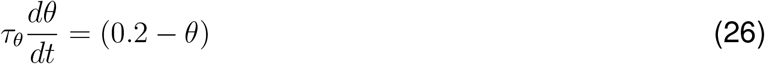

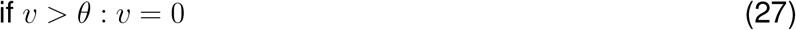

where *I*_*in*_ is the corresponding pixel value (On-population) or the pixel value multiplied by −1 (Off-population).

This experiment also adds an adaptive threshold to the neuronal model.

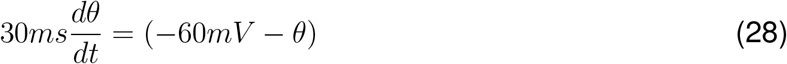

with *θ*(0) = −69*mV*. This increases the activity during the first stimuli presentations. Furthermore, the resting potential was set to −70mV and upon reset the membrane potential jumped to −75mV.

To characterise receptive fields, we compute the mean and standard deviation of the receptive field *r*_*i*_ of each excitatory neuron. To quantify the distribution of orientations of the receptive field of each network, we convolved each with differently oriented Gabor patches. The Gabor patches are defined as:

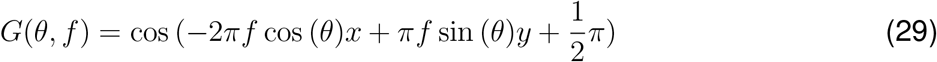

*x* and *y* are the pixel values relative to centre of the image, (*x, z* ∈ (−5, 5)). *θ* is the orientation of the Gabor patch and *f* the spatial frequency. Orientations are linearly spaced from −90° to 90°. Where 0° denotes a vertical oriented Gabor patch.

### Reproducibility notice and code availability

All code is available at https://github.com/prubisch/inhibitory_plasticity_regulation_in_voltage_dependent_plasticity_models. Any simulations which include randomised variables have a set random seed. Following python packages were used for simulation and analysis: numpy (version 1.23.5)^49^, matplotlib (version 3.8.4)^50^, scikit-learn (version 1.2.2)^51^ and the publicly available BRIAN2 package (version 2.5.1)^52^.

## ACKNOWLEDGMENTS

We would like to thank the members of the “ Neurons and Systems” group for helpful discussions and comments. We would like to thank Isabel Cornacchia and the anonymous reviewers for the helpful comments.

## AUTHOR CONTRIBUTIONS

Conceptualization, P.R. and M.H.H.; methodology, P.R.; investigation, P.R.; writing. P.R., M.I.S., M.H.H.; supervision, M.H.H.

## Appendix

### Illustration of the Voltage-Dependent Pathway model

The VDP models the downstream processes of plasticity induction as a simplified pathway-dynamic which is activated by depolarisation of the postsynaptic membrane. Therefore, the overall weight update from a single pairing can only be determined after several milliseconds when the pathway activations for LTP and LTD have both returned to 0. It emphasizes the temporal competition between LTP and LTD pathway activation as the central mechanism determining the sign and amplitude of the synaptic change. The illustration provided in Figure 7 visualises the mechanism used in the VDP.

**Figure 7.**
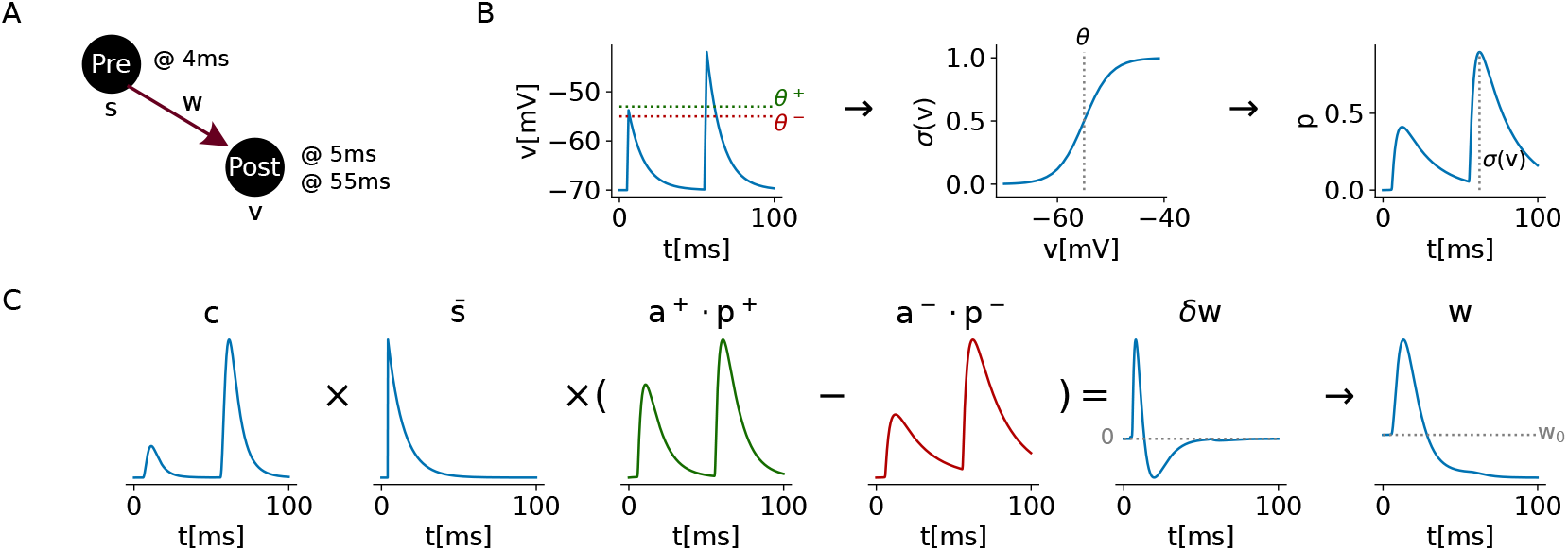
Illustration of the VDP model. **A**: Stimulation schema. The presynaptic spike train *s* and the postsynaptic membrane potential *v* are measured to compute the synaptic weight update. The time points denote spike and postsynaptic stimulation time to elicit the activation of the traces in B and C. **B**: Visualisation of the pathway induction process. The instantaneous membrane potential (left) is applied to a sigmoid function (middle) as a soft threshold with the middle points *θ*^+^ and *θ*^−^ for the potentiation and depression pathway. The result of the sigmoid operation is used as the maximum amplitude or offset for the biexponential pathway activation *p* (right). **C**: Illustration of the computation and functions in the synaptic weight update (equation 17). The component is a scaling factor mimicking calcium concentration dependent on the potentiation and depression pathway. This has been omitted in the equations to allow a direct comparison of the models. The scaling does not qualitatively influence the inhibitory effect. From left to right: The calcium level *c* is multiplied with the presynaptic trace 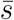 and the difference between the potentiation *p*^+^ and depression pathway *p*^−^ in order to calculate the momentary weight *w*. Due to the amplitude parameter *a*_+_ *<* 1, the illustrative example results overall in LTD, despite the weight increasing at first after the induction mechanism.

### Reproduction of central experimental results with the VDP

To test the validity of the VDP, we tested whether core results of the experimental literature focusing on excitatory synaptic plasticity can be reproduced. The results which show that the VDP can be considered a valid synaptic plasticity model are shown in Figure 8.

**Figure 8.**
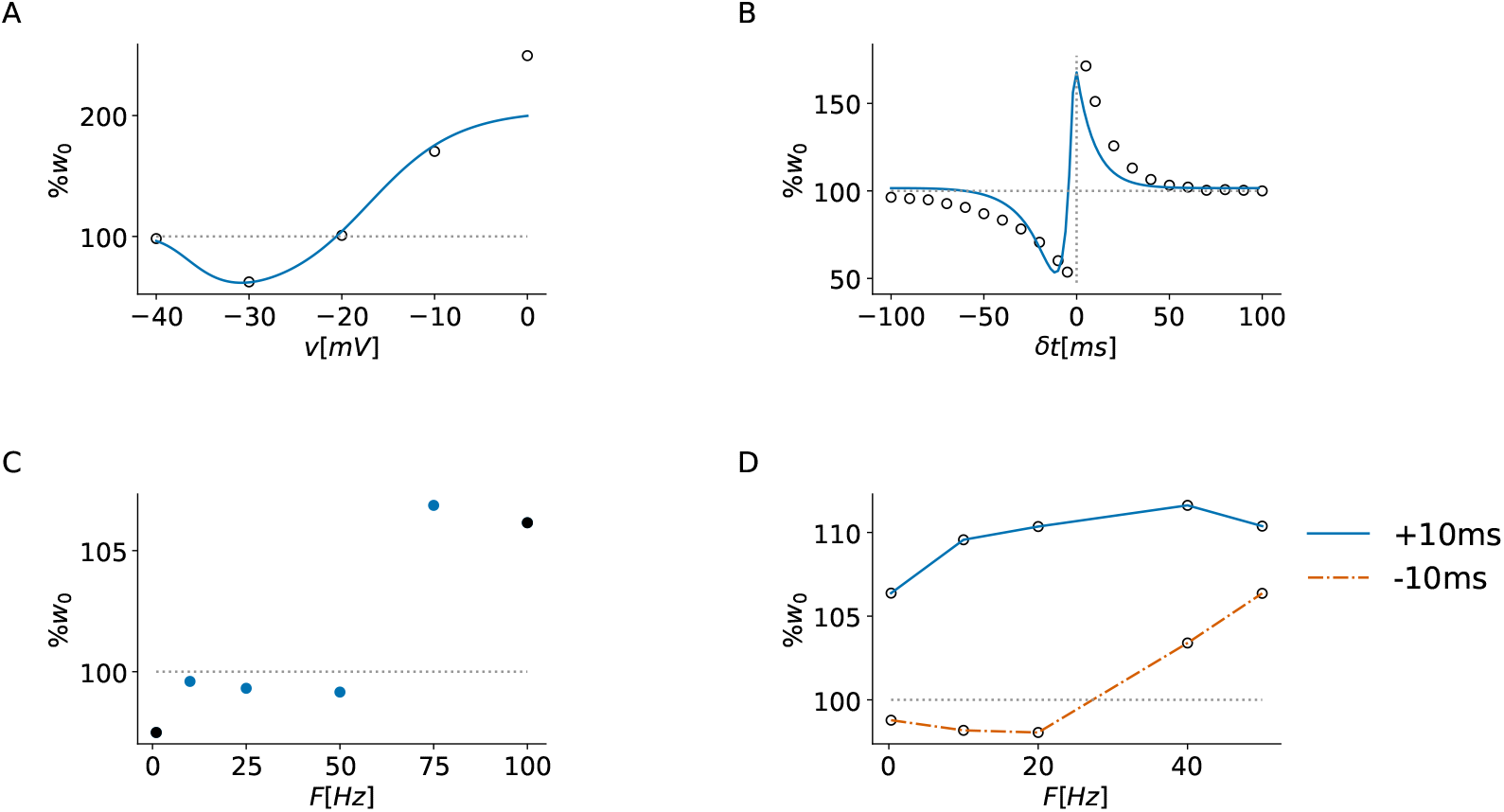
Predictions of the VDP for several experimental protocols. **A**: The voltage dependency follows the relationship as described by Ngezahayo et al. ^8^ closely. **B**: The VDP can reproduce the standard STDP kernel as measured by Froemke and Dan ^3^. The open-faced circles indicate the experimental measurements. **C**: Low Frequency Stimulation (LFS) and Theta Burst Stimulation effects (TBS) are reproduced. The solid circles indicate conditions measured in Dudek and Bear ^5^. **C**: The qualitative effect of the Pairing Frequency as described in Sjöström et al. ^2^ can be reproduced by the VDP. Open-faced circled here indicate the conditions tested by Sjöström et al. ^2^. They do not represent their measurements, as fitting the data quantitatively is limited by the description of the experimental protocol and neuron model used.

Furthermore, by changing the relative dynamics of the potentiation and depression pathway the VDP can reproduce different STDP kernel shapes as described in Abbott and Nelson ^1^. To characterise the different kernel shapes we measure the shape *s* defined as :

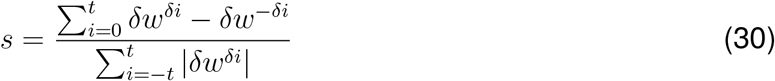

Figure 9 shows four distinct types of STDP kernel shapes alongside their *s* and *r* (LTP-LTD ratio) scores.

**Figure 9.**
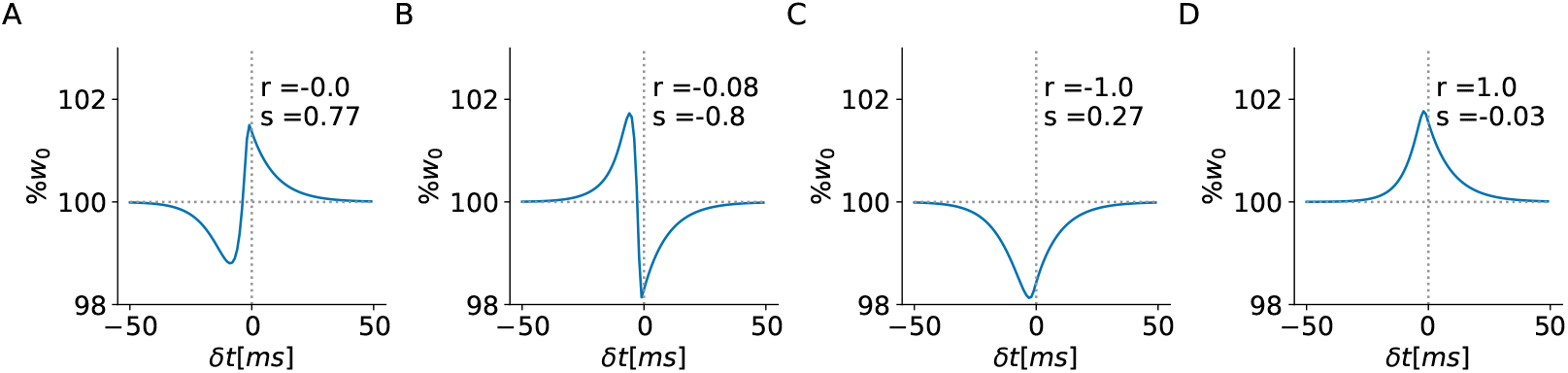
Different STDP kernel shapes reproduced by the VDP. The differences can be measured by calculating the kernel shape *s* and LTP-LTD ratio *r*. **A**: Balanced Hebbian STDP kernel with r = 0.0 and s = 0.77. **B**: Balanced anti-Hebbian STDP kernel with r = −0.08 and s = −0.8. **C**: LTD-dominated nearly-symmetric kernel. r = −1.0, s = 0.27. **D**: LTP-dominated nearly symmetric STDP kernel. r = 1.0 and s = −0.03.

### EI balance and membrane potential statistics in a Hodgkin-Huxley regular spiking neuron

All simulations use a LIF neuron as described in Section Methods. As this is a simplified neuron model without an explicit spike shape estimation, it is central to test whether the relationship between EI balance and membrane potential statistics holds in neuron models which approximate the spike shape. The results of the same simulation with a regular spiking Hodgin-Huxley model^53^(HH) are illustrated in Figure 10. It confirms that inhibitory-dominated synaptic input results in a higher mean membrane potential at similar firing rates in comparison to excitatory-dominated synaptic input. However, the dip in the variance of the membrane potential at low firing rates is not present in this simulation, indicating that this is an artefact caused by the (missing) approximation of the spike shape of the LIF neuron.

**Figure 10.**
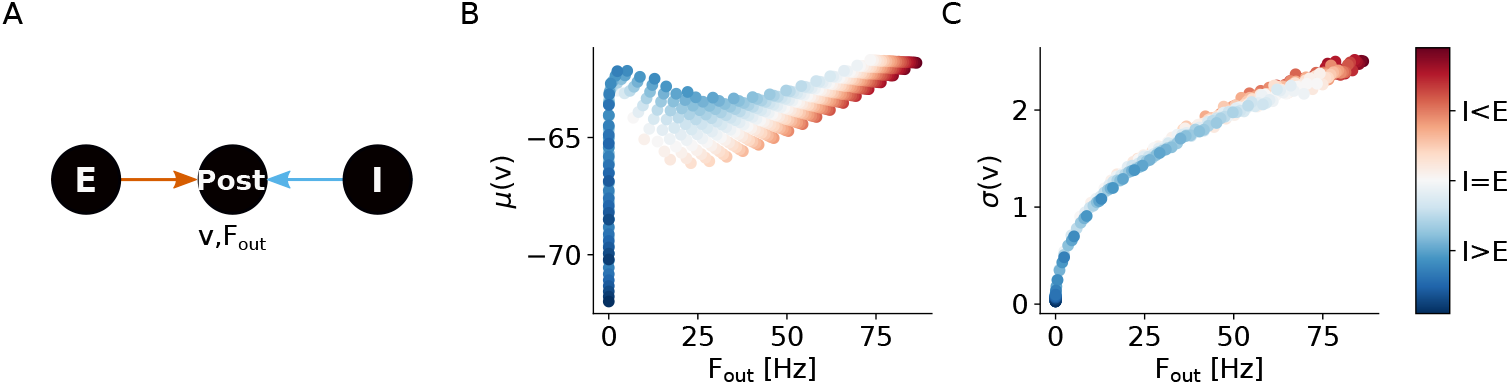
EI balance influences on mean and standard deviation of the membrane potential is similar in LIF neurons and HH neuron.**A** A postsynaptic HH neuron is driven by 100 excitatory and 100 inhibitory Poisson neurons with firing rates *F*_*E*_, *F*_*I*_ ∈ [0, 100)Hz (5Hz resolution). The colour-coding of the data points indicates the EI balance of the input computed as 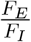. We measure the postsynaptic membrane potential, output frequency, as well as the weight change of an excitatory (orange,dashed) synapse. The presynaptic neuron of that synapse fires regularly with the same frequency as the excitatory Poisson population. All results are calculated over 10s simulation time. **B**: The mean membrane potential increases when the synaptic input is inhibition-dominated. **C**: The standard deviation of the membrane potential measuring the fluctuation amplitude is increasing when the synaptic input is dominated by excitation.

### Inhibitory frequency dependency in voltage-dependent plasticity models

The simulation results portrayed in Figure 3 also enable us to investigate the development of inhibitory weights. The presynaptic frequency is set to be equivalent to the frequency of the inhibitory Poisson population. The results for this simulation are shown in Figure 11. They show that the inhibitory weight development is strongly driven by the presynaptic firing rate. At high postsynaptic frequencies the update amplitude decreases progressively due to the decrease in presynaptic frequency. The models differ only slightly in their inhibitory frequency dependency and show similar qualitative behaviour.

**Figure 11.**
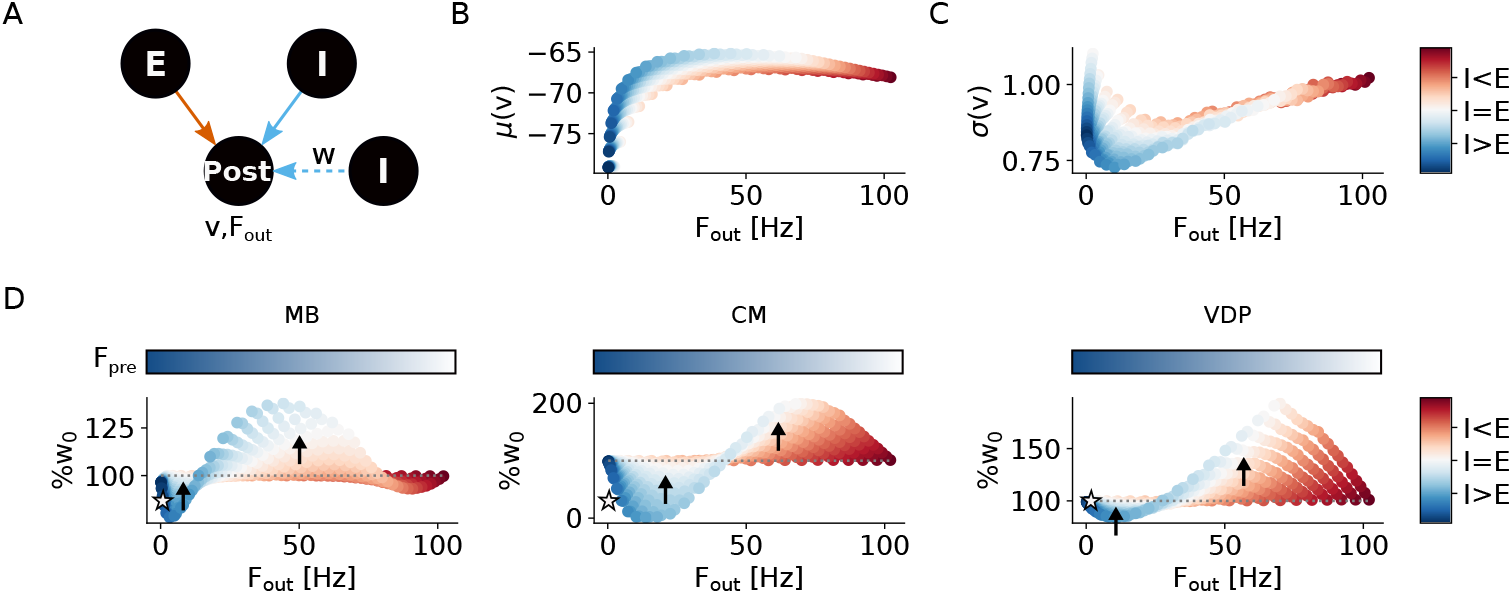
The frequency dependency of inhibitory synapses is driven by the presynaptic activity leading to similar patterns in all plasticity models. **A** A postsynaptic neuron is driven by 100 excitatory and 100 inhibitory Poisson neurons with firing rates *F*_*E*_, *F*_*I*_ ∈ [0, 100)Hz (5Hz resolution). The colour-coding of the data points indicates the EI balance of the input computed as 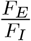. We measure the postsynaptic membrane potential, output frequency, as well as the weight change of an inhibitory (blue,dashed) synapse. The presynaptic neuron of that synapse fires regularly with the same frequency as the excitatory Poisson population. All results are calculated over 10s simulation time. **B**: The mean membrane potential increases when the synaptic input is inhibition-dominated. **C**: The standard deviation of the membrane potential measuring the fluctuation amplitude is increasing when the synaptic input is dominated by excitation. **D**: Plasticity predictions of the plasticity models of the inhibitory weights. Model identity is indicated by the abbreviations on top. All results are averaged over 100 simulation runs. The color bar on top indicates the frequency of the regular spiking presynaptic neuron. The open-faced star indicates the regime of pathological behaviour at low postsynaptic frequencies (negative plasticity-activity feedback). Arrows indicate the direction of EI balance shift due to the predicted synaptic plasticity update.

### Filter properties of the plasticity models

The postsynaptic filter of the MB and CM are low-pass filters by definition. However, adding the threshold and in case of the CM combining a filtered and an unfiltered condition in the plasticity induction introduces conditional filtering. The VDPs down-stream filter in form of the pathway activities can neither be described as a low-pass nor high-pass filter by definition. Therefore, we simulated a burst of 5 spikes (200Hz) repeated with either 5Hz (Figure 12a) or 50Hz (Figure 12b) to visualise the differences in the filter properties of the models.

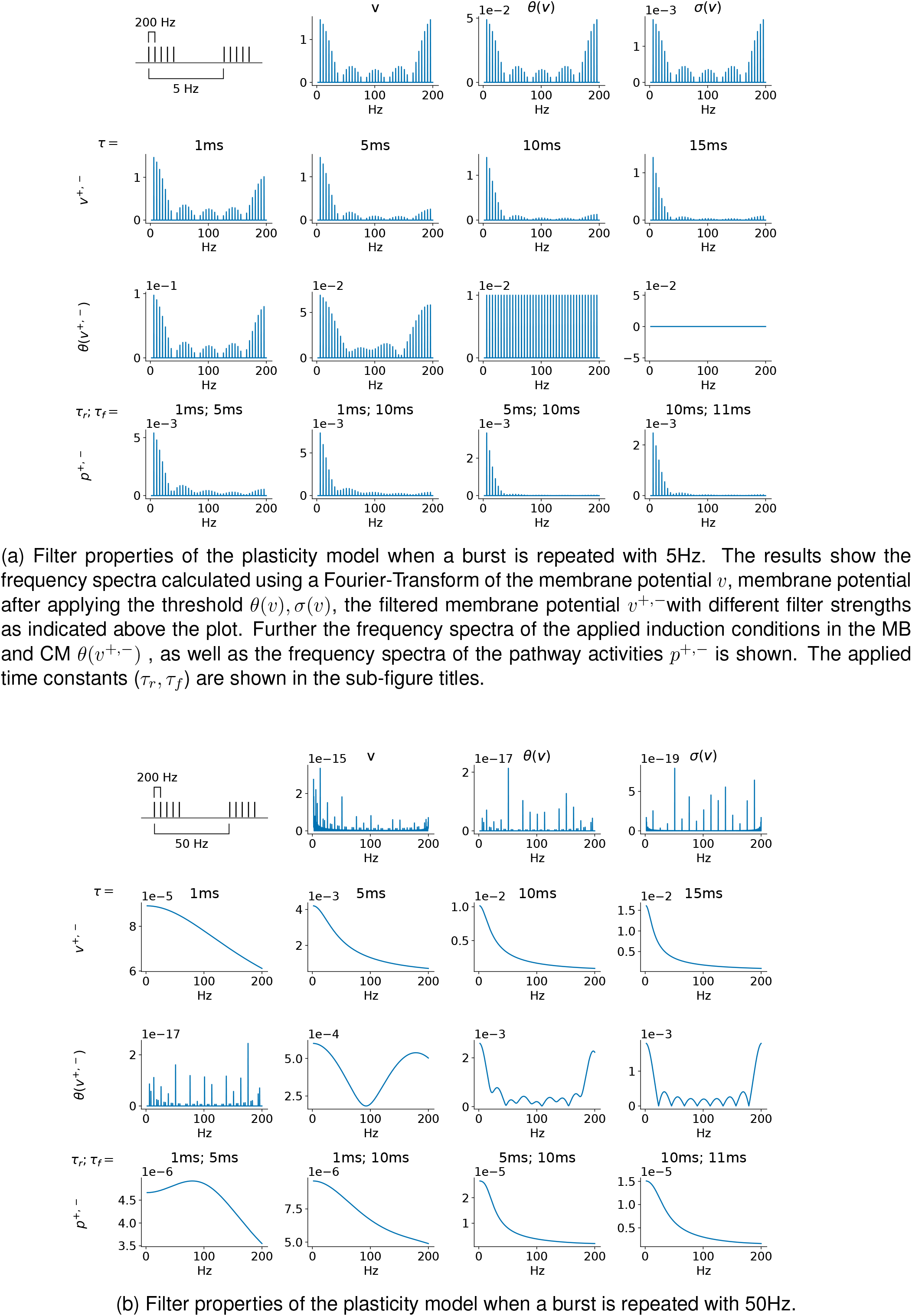

### Influence of time constants on the sensitivity to inhibition onset

A central question which arises, when comparing different synaptic plasticity models is whether similar results can be achieved simply by adapting model parameters. In this work we argue that the temporal extent of the inhibitory regulation as well as the sensitivity is dependent on the model definition. In the MB and CM models show a parameter-sensitive fast fluctuation sensitivity, while the VDP’s fast fluctuation sensitivity is a model feature and independent of its parametrisaiton. To demonstrate the parameter influence on the plasticity patterns elicit by changing stimuli, we vary the time constants and measure how they influence the slope of weight change of the mechanistic simulation illustrated in Figure 4. Figure 13 demonstrates that the presynaptic temporal constant has the biggest effect in the MB anc CM model behaviour by allowing a longer effective pairing window in these models. The VDP has slower postsynaptic dynamics which is why the presynaptic constant has a less strong effect in this model while the postsynaptic constants influence the slope by prolonging pathway activity.

**Figure 13.**
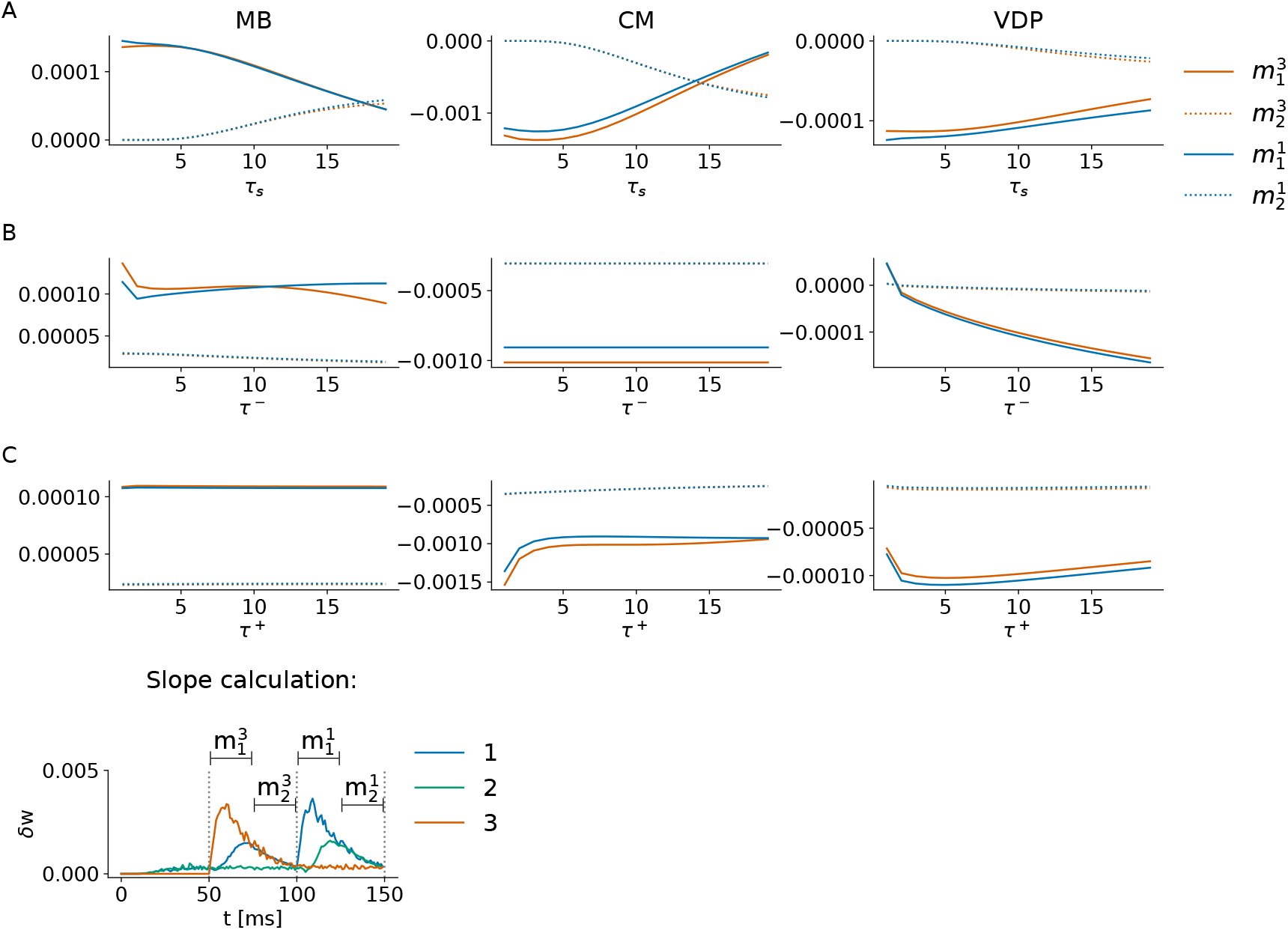
Different STDP kernel shapes reproduced by the VDP. The differences can be measured by calculating the kernel shape *s* and LTP-LTD ratio *r*. **A**: Balanced Hebbian STDP kernel with r = 0.0 and s = 0.77. **B**: Balanced anti-Hebbian STDP kernel with r = −0.08 and s = −0.8. **C**: LTD-dominated nearly-symmetric kernel. r = −1.0, s = 0.27. **D**: LTP-dominated nearly symmetric STDP kernel. r = 1.0 and s = −0.03.

### Within-network receptive field similarity

Figure 14A shows further results on the clustering of the receptive field statistics. To investigate the robustness of the clusters, the statistics were randomly shuffled (grey data). The similarity between receptive fields of the same network and its relationship to the variance of the receptive fields, Figure 14B-D, show the gradual change of receptive fields in the CM and VDP network, while the MB network builds clusters of receptive field with similar structures which is also captured by the clustering in their descriptive statistics.

**Figure 14.**
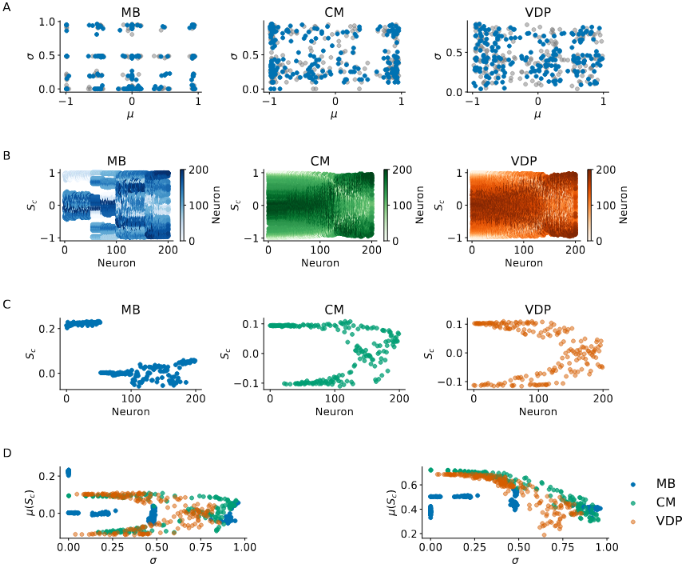
Clustering and further similarity analysis of the receptive fields. **A**:Original pairings of mean *µ* and variance *σ* (blue) and reshuffled pairs (gray). **B**: Cosine similarity *S*_*c*_(*i, j*) of the neurons *i* and *j* receptive field. **C**: Mean cosine similarity *µ*(*S*_*c*_(*i, j*)) or neuron *i*. **D**: Relationship between the standard deviation *σ* and *µ*(*S*_*c*_(*i, j*)) (left) and *µ*|(*S*_*c*_(*i, j*))| (right).

### Parametrisation details

**Table 2:**
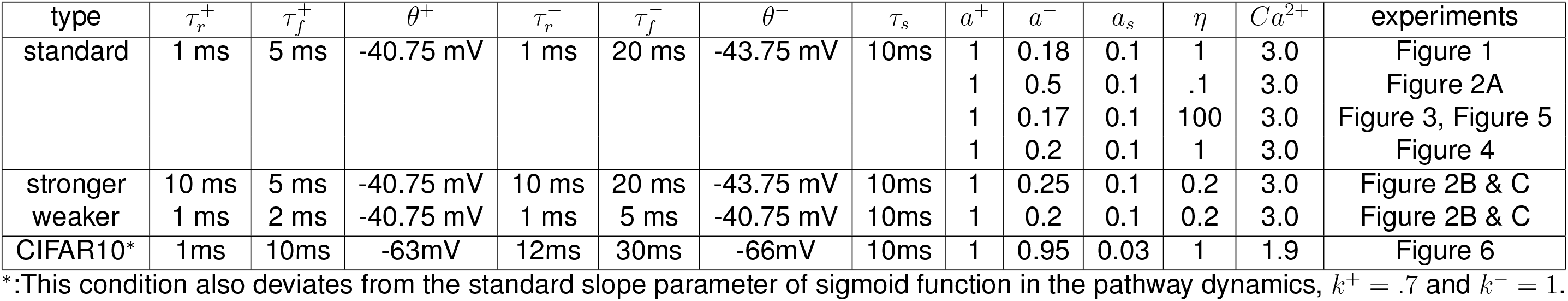
Parametrisation of the VDP.

**Table 3:**
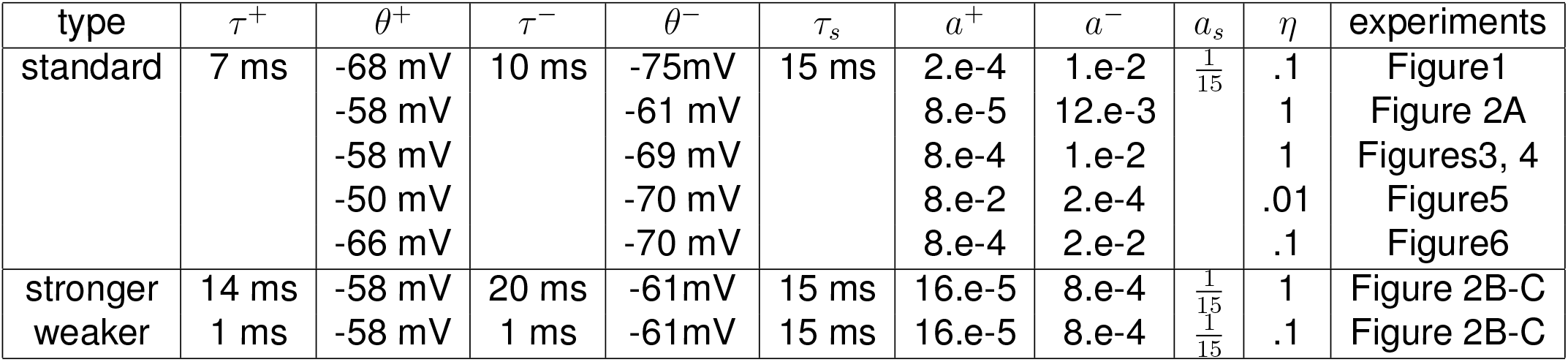
Parametrisation of the CM.

**Table 4:**
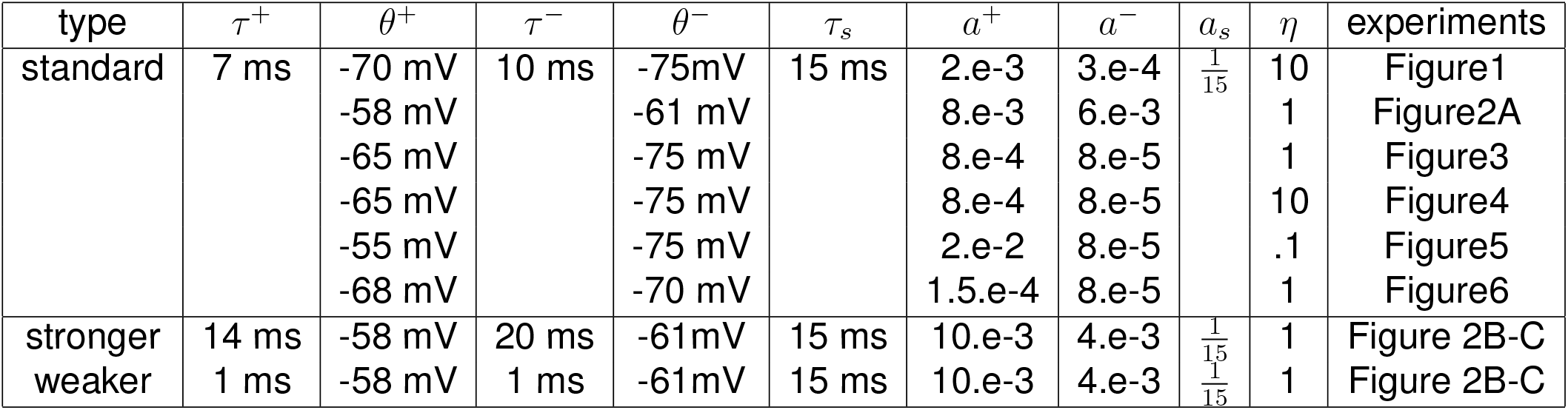
Parametrisation of the MB.

